# The role of genus and life span in predicting seed and vegetative trait variation and correlation in *Lathyrus*, *Phaseolus*, and *Vicia* (Fabaceae)

**DOI:** 10.1101/2021.02.17.431656

**Authors:** Sterling A. Herron, Matthew J. Rubin, Matthew A. Albrecht, Quinn G. Long, Marissa C. Sandoval, Allison J. Miller

## Abstract

**PREMISE OF THE STUDY:** Annual and perennial life history transitions are abundant among angiosperms, and understanding the phenotypic variation underlying life span shifts is a key endeavor of plant evolutionary biology. Comparative analyses of trait variation and correlation networks among annual and perennial plants is increasingly important as new perennial crops are being developed in a predominately annual-based agricultural setting. However, it remains unclear how seed to vegetative growth trait relationships may correlate with life span.

**METHODS:** We measured 29 annual and perennial congeneric species of three herbaceous legume genera (*Lathyrus*, *Phaseolus*, and *Vicia*) for seed size and shape, germination proportion, and early vegetative height and leaf growth over three months in order to assess relative roles of genus and life span in predicting phenotypic variation and correlation.

**KEY RESULTS:** Genus was the greatest predictor of seed size and shape variation, while life span consistently predicted static vegetative growth traits. Correlation networks revealed that annual species had significant associations between seed traits and vegetative traits, while perennials had no significant seed-vegetative associations. Each genus also differed in the extent of integration between seed and vegetative traits, as well as within-vegetative trait correlation patterns.

**CONCLUSIONS:** Genus and life span were important for predicting aspects of early life stage phenotypic variation and trait relationships. Differences in phenotypic correlation may indicate selection on seed size traits will impact vegetative growth differently depending on life span, which has important implications for nascent perennial breeding programs.

---

Life history strategy in plants involves complex patterns of reproductive, growth, and survival trait allocation and trade-offs, which can shed light on past adaptive drivers and influence future evolutionary trajectories (Stearns, 1992). Describing ecological, genetic, and phenotypic life history patterns and commonalities among diverse taxa is thus fundamental in advancing evolutionary biology (Díaz et al., 2016; Friedman and Rubin, 2015). Annuals and perennials are two broadly recognized life span categories in plants (Friedman, 2020). While annual plants grow, reproduce, and senesce within a single year, perennial plants vary from small, short-lived herbaceous individuals that live 3-5 years to expansive, ancient clonal colonies (e.g., *Populus*) and woody individuals (e.g., *Sequoia*) that can live for hundreds or thousands of years. Although traits associated with annual-perennial differences are well-defined in some model systems, this comparative framework has garnered renewed attention with increasing focus on understudied, wild species as sources of novel crop breeding material, particularly perennial species closely related to annual crops, such as corn, rice, and wheat (Lundgren and Des Marais, 2020). Additional analyses across diverse, non-model systems are needed to determine the phenotypic and genetic factors underlying differences in annual and perennial species (Friedman, 2020).

Several broad life history frameworks have been proposed to interpret the large range of phenotypic variation and correlation in plants. Classic plant life history theory balances an individual’s survival with reproduction as modes of achieving fitness (Cole, 1954; Charnov and Schaffer, 1973; Gadgil and Solbrig, 1972). Derivations of this have described systems of interconnected phenotypes which span predictable trait networks or spectra (e.g., Grime, 1977; Diaz et al., 2016). One example, the plant economic spectrum, categorizes plants by their resource-use strategy: resource-acquisitive plants use available resources for immediate structural growth, and resource-conservative plants invest resources in expensive storage organs to prioritize future growth and survival (Chapin, 1980; Chapin et al., 1990; Reich, 2014). The plant economic spectrum has been primarily studied in leaves, with acquisitive plants having broad, short-lived leaves and a high photosynthetic rate, and with conservative plants having thick, long-lived leaves and a lower photosynthetic rate (Wright et al., 2004). This economic framework has also been extended to stems (Chave et al., 2009) and roots (Roumet etl al., 2016).

Within the framework of the plant economic spectrum, annuals are generally categorized as acquisitive species and perennials as conservative species. Evidence for differences in traits associated with life span comes largely from leaf and shoot traits (e.g., specific leaf area, leaf water content, and relative growth rate are greater in annuals than perennials; Garnier, 1992; Garnier and Laurent, 1994; Atkinson et al., 2016), reproductive traits (e.g., seed mass fraction and reproductive energy percent allocation are greater in annuals; Pitelka, 1977; Vico et al., 2016) and root traits (e.g., specific root length and root nitrogen concentration are greater in annuals, root tissue density and root diameter are greater in perennials; Roumet et al., 2006). Nevertheless, it has long been recognized that plant species of different life histories are capable of having diverse trait systems that are not always predicted by generalized models; these traits are ultimately determined by the species’ distinct environmental stressors and evolutionary history (Thompson and Hodkinson, 1998; Crews and DeHaan, 2015). Herbaceous perennials as a group have been recognized as particularly variable in life history trait combinations (Grime, 1977; Kitchen, 1994). Several studies have found evidence for wild herbaceous perennial species with acquisitive and ruderal traits, including comparable reproductive biomass and relative growth rate to annuals (Verboom et al., 2004; González-Paleo and Ravetta, 2015; Vico et al., 2016).

In order to understand evolutionary patterns of life history strategy, it is helpful to study phenotypic variation associated with life span within and among different phylogenetic lineages. Life span transitions are common in many angiosperm systems (e.g., Castillejinae: Tank and Olmstead, 2008; core Pooideae: Lindberg et al., 2020; Saxifragales: Soltis et al., 2013), which allows for investigation into phenotypic and genetic commonalities in different life spans. Previous studies in grasses and other systems have found similar annual-perennial phenotypic differences across multiple genera, including in shoot, leaf, root, and reproductive traits (as discussed above; Garnier, 1992; Garnier and Laurent, 1994; Garnier and Vancaeyzeele, 1994; Vico et al., 2016), and accounting for phylogenetic relationship of the studied taxa has shown to be key in discerning life span differences apart from other evolutionary influences (Garnier, 1992). However, the phenotypic and genetic factors underlying life history and life span variation remain uncharacterized in many diverse plant groups.

Networks of trait correlation are also fundamental to studies of life history diversity. Determining relationships between plant organs, or phenotypic integration, helps to conclude whether key traits were consistently selected together as a functional module, if trade-offs are occurring, or if the traits in question remain relatively independent (Murren, 2002). These trait relationships can clarify fundamental constraints on plant phenotypic diversity. Previous studies in herbaceous species have found negative relationships between vegetative growth and some aspects of reproduction (e.g., earlier flowering; Geber, 1990) but generally positive relationships between vegetative size traits and size of seeds produced within a species (Geber, 1990) and among species (Kleyer et al., 2019). Other studies have further found that life span plays an important role in predicting functional reproductive-vegetative trait relationships, including the association of flowering time with seed size and plant height, with seed size being derived from separate studies (Bolmgren and Cowen, 2008; Du and Qi, 2010) or the size of seeds produced by the plants on which vegetative traits were measured (Segrestin et al., 2020), and between leaf area and seed size (Hodgson et al., 2017). However, to our knowledge few studies have focused in detail on congeneric annual-perennial differences in seed size vs. vegetative plant size / growth rate correlations. Here we seek to build on previous life history studies by investigating specifically how relationships between seed and vegetative traits in multiple genera may shift with life span.

Due to its economic and ecological importance, as well as biological diversity, the legume family (Fabaceae Lindl.) is an excellent system in which to study phenotypic patterns associated with life span. Specifically, subfamily Papilionoideae DC. contains numerous genera of agricultural importance with annual and herbaceous perennial congeners (Ciotir et al., 2019). Similar to other systems, annual legume species tend to have greater relative allocation to sexual reproduction than perennial congeners in terms of percent energy allocation (Pitelka, 1977) and percent dry biomass allocation (Turkington and Cavers, 1978). Studies have also explored other functionally important traits in congeneric annual and perennial legumes, including seed mass (Pitelka, 1977; Marshall et al., 1985; Kelly and Hanley, 2005; Ward et al., 2011; Herron et al., 2020); specific leaf area (den Dubbelden and Verburg, 1996; Roumet et al., 2000); photosynthetic rate (Pitelka, 1977; Roumet et al., 2000); and growth rate (den Dubbelden and Verburg, 1996; Kelly and Hanley, 2005). In these studies, life span was commonly not the central focus and results were mixed, or there were equivocal patterns in terms of whether annuals and perennials showed greater trait values. These studies also tend to focus on two or three species per genus, while a much greater range of diversity exists for both annuals and perennials within genera and across Fabaceae (Ciotir et al., 2019).

Exploring patterns of life span-associated differences in seeds and vegetative traits will be valuable not only for better understanding their evolutionary background but also for informing crop breeding. Although herbaceous perennial species were generally not domesticated by early farmers (Van Tassel et al., 2010), attention is now focusing on these species as a potential means to slow or reverse soil erosion associated with annual grain agriculture and to sustainably yield in marginal environments (Glover et al., 2010; Ryan et al., 2018). An increasing number of diverse crop wild relatives are being considered for trait conferral via introgression or *de novo* domestication, including herbaceous perennial grains and pulses. Capturing the range of genetic variation available in wild germplasm will be essential for guiding breeding decisions (Schlautman et al., 2018; Smýkal et al., 2018). Characterizing phenotypic correlation within plants may form a predictive foundation for early-stage selection based on seeds, and will additionally inform whether breeding for certain phenotypes may lead to trade-offs with other desired traits, e.g., resource competition between reproductive and vegetative organs such as roots (González-Paleo et al., 2016; Pastor-Pastor et al., 2018).

Here we focus on a panel of annual and herbaceous perennial species from the agriculturally important Fabaceae genera *Lathyrus* L., *Phaseolus* L., and *Vicia* L. in order to better characterize phenotypic diversity associated with life span and add to our understanding of trait variation and correlation associated with different life history strategies. Starting with seeds, we measured phenotypic variation in early life stages: seed size and shape, germination, and seedling vegetative growth. We ask the following questions: 1) What is the relative importance of genus and life span in predicting seed and vegetative trait variation? 2) How do seed and vegetative trait correlations differ across genera and life spans? We address these questions by quantifying seed size, germination, and early life stage vegetative growth traits in congeneric wild, annual and perennial species grown in a common environment.

## MATERIALS AND METHODS

### Plant material

We selected annual and herbaceous perennial species from three economically important legume genera: *Lathyrus* (grass pea), *Phaseolus* (common bean), and *Vicia* (vetch). We obtained 80 accessions (Appendix S1) from the United States Department of Agriculture’s National Plant Germplasm System (Western Regional Plant Introduction Station, Pullman, Washington, USA) in spring 2017, which were stored in a desiccator at 3 to 4.5°C, 36% relative humidity prior to the start of the experiment. All accessions used were designated as “wild” by the USDA (with the exception of one accession of *V. americana*, see below), meaning that seed collection took place in a natural population outside of cultivation, although this designation may include feral and/or naturalized escapees from cultivation in the case of some species collected outside their native range (primarily *Lathyrus* collected in the U.S.; Kenicer, 2008). All accessions were harvested from a seed increase effort at the germplasm center, except for one accession of *Lathyrus japonicus* (W6 45319) and *L. latifolius* (PI 602368), which were collections made directly from the original wild population. Time of seed increase / collection ranged from two to 30 years prior to this study, the duration of which seeds were kept in frozen storage. For this experiment, accessions were germinated and then grown from June to September 2017. For *Lathyrus*, five annual and five perennial species were studied; for *Phaseolus*, four annual species and five perennial species; and for *Vicia*, five annual species and five perennial species (Table 1). One to seven accessions were obtained for each species, with origins from diverse geographic areas when possible. All accession information is available in Appendix S1 (see Supplemental Data with this article).

**Table 1.**
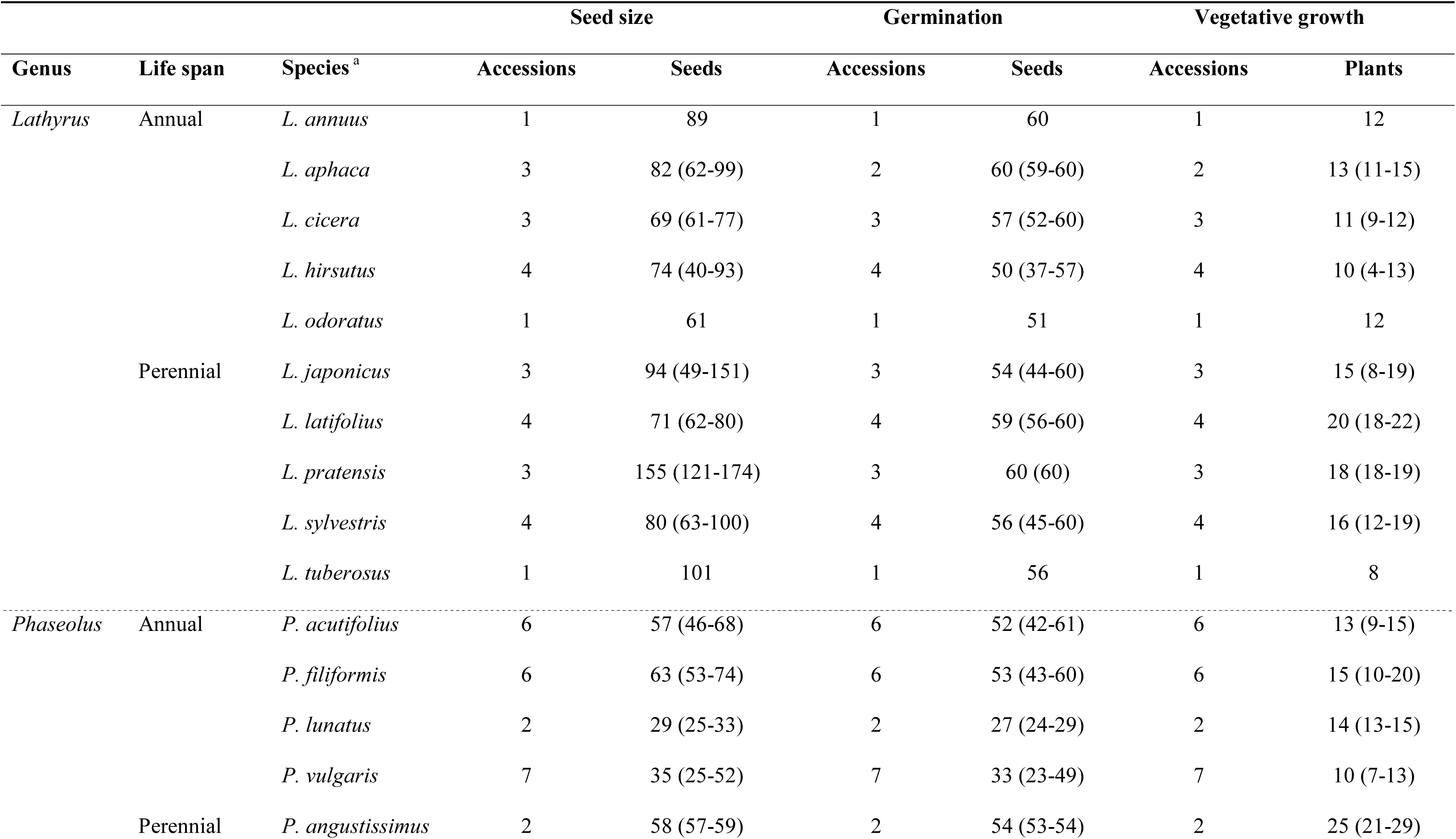
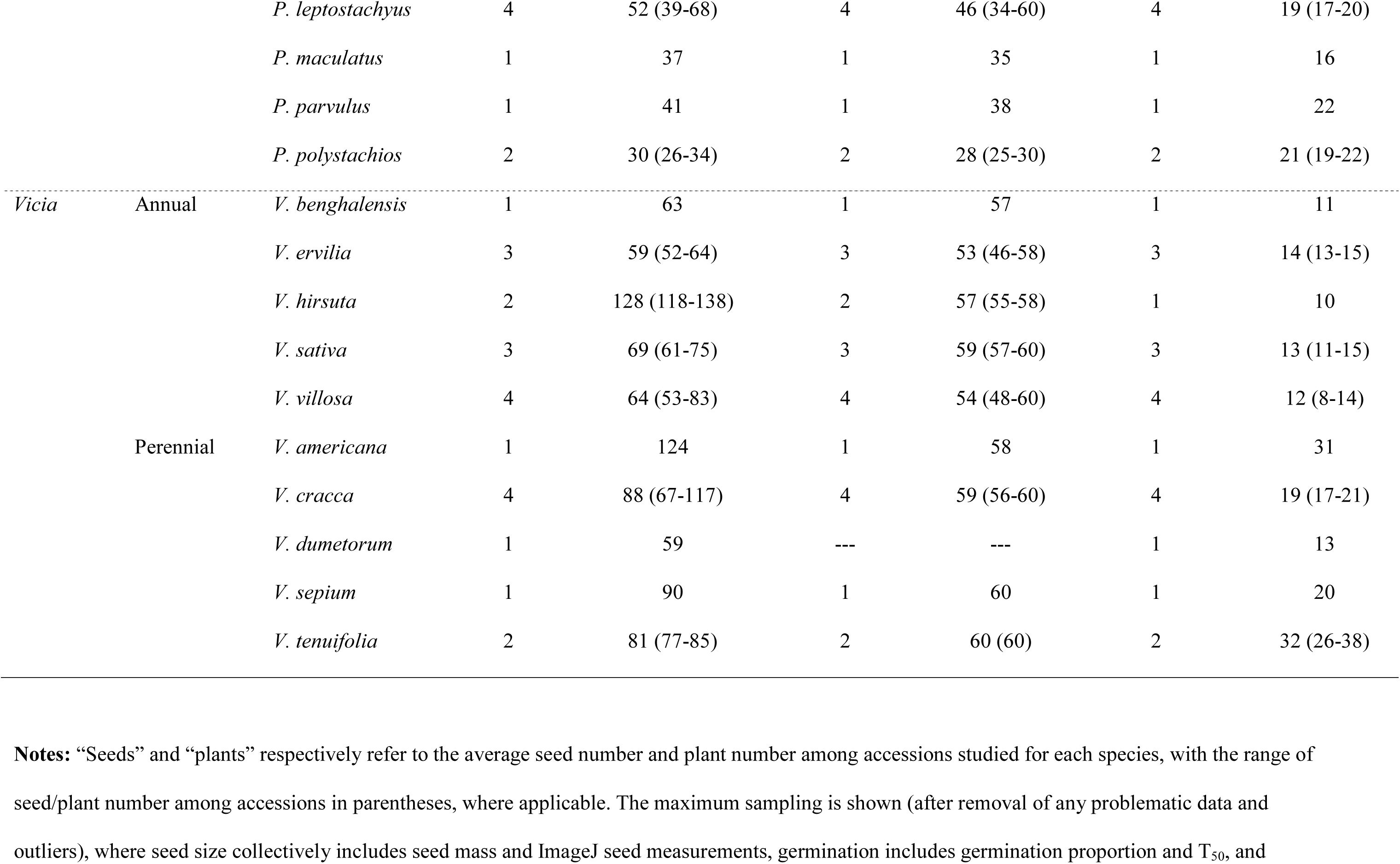
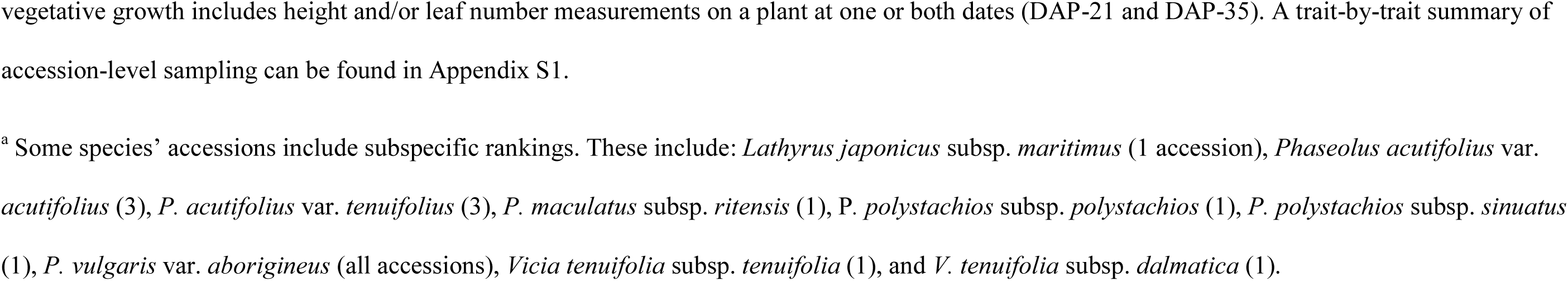
Sampling summary of seed, germination, and vegetative traits in annual and perennial species of *Lathyrus, Phaseolus,* and *Vicia*.

When possible, we obtained accessions of annual and perennial species from closely related phylogenetic groups based on the most recently published, comprehensive molecular phylogenies of each genus. *Lathyrus* and *Vicia* are both within the economically important tribe Fabeae, which also includes pea, lentil, and grass pea (Schaefer et al., 2012). Both genera consist of c. 160 herbaceous annual and perennial species distributed globally (Lewis et al., 2005). Within *Lathyrus*, clades sampled here include: 1) the clade including sections Aphaca (*L. aphaca*), Pratensis (*L. pratensis*), and Orobus (*L. japonicus*); and 2) the large section Lathyrus (*L. annuus*, *L. cicera*, *L. hirsutus*, *L. odoratus*, *L. latifolius*, *L. sylvestris*, and *L. tuberosus*) (Asmussen and Liston, 1998; Schaefer et al., 2012; Table 1). Within *Vicia*, clades sampled here included 1) section Pedunculatae (*V. americana*, *V. dumetorum*); 2) section Ervilia (*V. ervilia*, *V. hirsuta*); 3) section Vicia (*V. sepium* and *V. sativa*); and 4) section Cracca (*V. cracca*, *V. tenuifolia*, *V. villosa*, and *V. benghalensis*) (Endo et al., 2008; Castiglione et al., 2011; Schaefer et al., 2012; Jaaska, 2015; Table 1). Notably, section Ervilia is more distantly related to other sections of *Vicia*; it is sister to all clades of the tribe Fabeae (including *Vicia* and *Lathyrus*, among other genera), and it has been proposed that it be raised to genus level (Schaefer et al., 2012). These changes have yet to be finalized, so we retain the current taxonomy of section Ervilia. *Phaseolus*, a monophyletic genus of c. 70-80 species, is more distantly related to *Lathyrus* and *Vicia* and is in tribe Phaseoleae (Freytag and Debouck, 2002; Delgado-Salinas et al., 2006). For *Phaseolus*, two broad clade groups were sampled here: the 1) the Filiformis group (*P. angustissimus* and *P. filiformis*) and the Vulgaris group (*P. acutifolius* and *P. vulgaris*); and 2) the Polystachios group (*P. maculatus* and *P. polystachios*), the Lunatus group (*P. lunatus*), and the Leptostachyus group (*P. leptostachyus*) (Delgado-Salinas et al., 2006; Table 1). Also included is the more distantly related *P. parvulus* in the Pauciflorus group (Delgado-Salinas et al., 2006; Table 1). The USDA has since updated the taxonomic status of *P. maculatus* subsp. *ritensis* to *P. ritensis*, but we retain its original taxonomic status here (Freytag and Debouck, 2002). The *Lathyrus* species in this study are primarily native to temperate and subtropical regions of Eurasia, with *L. japonicus* having a global temperate distribution (Wu et al. 2010; Schaefer et al., 2012; Global Biodiversity Information Facility; http://www.gbif.org). *Vicia* species in this study are native to temperate regions of Eurasia (though some also extend to subtropical regions), with the exception of the temperate North American species *V. americana* (Wu et al. 2010; Schaefer et al., 2012; GBIF.org). Most *Phaseolus* species here are from arid climates in the southwestern U.S. into northwest Mexico, or are native to dry or moist habitats in Mexico, Central America and South America, with *P. polystachios* standing out as the only *Phaseolus* species whose native range extends into temperate regions (Freytag and Debouck, 2002; Bitocchi et al. 2017; GBIF.org).

For each accession used in this study life span and cultivation status were derived from the online USDA description in GRIN-Global (Germplasm Resources Information Network) and confirmed with literature (Wu et al. 2010; Schaefer et al., 2012; Freytag & Debouck 2002). We use the annual and perennial life span classifications only as a preliminary framework for understanding trait variation, acknowledging that life span in the strict sense is only part of a broader, complex life history strategy. In the cases where the USDA life span assignment (annual or perennial) and literature were contradictory, we conducted extensive literature research and expert consultation (Daniel Debouck, pers. comm., 2020). This occurred for *Phaseolus filiformis*, a Southwestern U.S. desert ephemeral (annual) (Buhrow, 1983; Nabhan and Felger, 1985; Freytag and Debouck, 2002), and *P. lunatus*, a species with an expansive distribution from Mexico to South America and which is variable in life span depending on the minimum temperature and the severity of the dry season where it is growing (Freytag and Debouck, 2002; Bitocchi et al., 2017). Although *P. lunatus* is capable of perenniality in mild, aseasonal climates, unlike other perennials in this study its fibrous root system lacks substantial reserves and is not able to endure extended harsh conditions (Freytag and Debouck, 2002); thus, in the absence of specific life span information for source populations we deferred to the USDA’s original assignment of annual. Due to life span ambiguity, the linear models were also tested with *P. lunatus* accessions removed to confirm that it was not driving any trends (see below). If there was uncertainty regarding the cultivation status or origin of an accession, the original collection information was consulted and collectors contacted when possible to confirm our information. For example, the accession of *V. americana* from Canada (PI 452486) was noted as “cultivated” in the USDA system; while it may sometimes be seeded in restoration efforts, *V. americana* is not known to be domesticated and this accession’s likely wild status was confirmed by the Plant Gene Resources of Canada, which also curates the accession.

### Seed size and shape measurements

Seed mass and two-dimensional seed shape parameters were characterized for each accession prior to sowing. A pool of seeds for each accession was first weighed on an Ohaus^®^ Adventurer^®^ Pro precision balance (Parsippany, New Jersey, USA) to the nearest 0.1 mg and divided by the total number of seeds for that accession to estimate mean single seed mass. Small funicles (rare) were not removed before weighing, except in the case of *Vicia sepium*, where uniquely conspicuous funicles were present on a few seeds. Seeds were then scanned at a resolution of 1200 dpi using an EPSON DS-50000 scanner (Nagano, Japan). When seeds were bilaterally symmetric (in *Lathyrus cicera* and all *Phaseolus*), they were oriented with the flat (lateral) side facing down and the hilum parallel lengthwise to the scanner surface. After scanning, images were converted from a CMYK jpg to png (400dpi) and analyzed in ImageJ (Schneider et al., 2012). Severely damaged seeds and seeds in a non-standardized orientation (only occurring for a few bilaterally symmetric seeds of *Lathyrus cicera*) were removed from the analysis. In ImageJ seed images were cropped and converted to binary (“Make Binary” function), or if the contrast was not well defined, the image was converted to 8-bit grayscale and a binary threshold (“Threshold” function) was applied and the threshold value adjusted for the highest seed contrast and lowest noise (shadows). Remaining pixel holes within seeds were removed using the “Fill Holes” function, and erroneous gaps on the perimeter of seeds were filled using the “Convex Hull” function (gift wrapping algorithm) or traced in manually when this was not adequate. Accessory structures still attached to the seeds (primarily funicles) were manually removed from the image. From this final image we extracted four size parameters per seed: length (Feret’s diameter, or the maximum distance between any two points along the perimeter of the seed), width (minimum distance between any two points along the perimeter of the seed), perimeter, and area, and two shape parameters, circularity and roundness. Circularity is calculated as 4π × (Area)/(Perimeter), and represents the extent to which the seed shape approximates a circle, ranging from 0 (infinitely elongated shape) to 1 (perfect circle). _Roundness is calculated as 4 × (Area)/(_π × (Major axis length)^2^) and is the inverse of the seed’s aspect ratio (length to width ratio of the best fitting ellipse); as this value approaches 1 it represents a width that is closer to the length of the seed. A majority of the seeds measured for size and shape were randomly chosen for use in germination experiments. In this study, seed traits are the initial juvenile stage in plants later measured for vegetative growth.

### Germination measurements

All seeds were sterilized in 6% sodium hypochlorite aqueous solution for 5 to 6.5 minutes, then rinsed with water purified via reverse osmosis (RO water) and patted dry (Frehner and Conn, 1987; Galasso et al., 1997). Due to the prevalence of physical dormancy induced by a water-impermeable seed coat in legumes (Baskin and Baskin, 2014), all seeds were scarified collectively by accession with sandpaper, pressing firmly until a breach of the seed coat was visible on the majority of seeds; this was completed in short (∼10 sec) bursts, usually 3-5 times. This was done without specificity to any part of the seed scarified, to simulate heterogeneous, natural scarification. P100 or P60 grade sandpaper was used depending on the observed hardness of the seed coat. A subset of seeds was taken randomly from each accession for germination, excluding malformed and broken seeds. Minimal damage occurred from the germination protocol, except for one accession of *Vicia hirsuta* (PI 219631) where about half of the seeds were damaged from bleach sterilization, which were not used. Germination models were tested with this accession removed to determine any impact (see sect. 2.3). All scarified seeds were surface-sown in unsterilized quartz sand (Fairmount Santrol Handy Sand (Chesterland, Ohio, USA); approximately 34 mL) in 20 mm deep plastic petri dishes; seeds were oriented horizontally (morphology allowing, hilum parallel lengthwise to substrate surface) in a grid pattern. Dishes were initially watered to saturation (∼11-12 mL) and were remoistened after one week and subsequently when dry. Excess water and condensation were blotted off. All seeds were germinated in 12:12 h light:dark conditions inside a temperature-controlled incubator, with 20:10°C and 25:15°C light:dark temperature settings for temperate (“temperate settings”) and subtropical / tropical species (“tropical settings”), respectively, in order to expose the species to temperatures similar to their native range. *Vicia* seeds were incubated at temperate settings, *Phaseolus* seeds were incubated at tropical settings, and *Lathyrus* seeds were divided into both settings depending on the species (*Lathyrus* annuals included both temperate and subtropical / Mediterranean species, perennials only temperate) (Appendix S1). Each accession had 2-3 replicate petri dishes and 5-26 seeds per replicate. Replicate dishes were randomized when placed in the incubator and re-randomized after each germination check. Germination was defined as an extension of the radicle past the seed coat, or in rare cases backwards emergence of the seedling due to the radicle pushing against the seed coat.

Germinated and imbibed seeds were counted beginning one day following placement on the substrate, then every one to four days up to 10-12 days, and then at three weeks and/or four weeks, or until all imbibed, viable seeds germinated (Baskin and Baskin, 2014). A subset of four to 38 (mean 15) vigorous seedlings were planted soon after germination, in order to have at least three separate replicates of five seedlings for each accession in most cases (see vegetative growth section). The remaining seedlings were returned to that accession’s petri dish; after this point, germination counts were based on the subtraction of ungerminated seeds from the original total seeds in the petri dish. Seeds which decayed and developed severe fungal infections were considered nonviable and removed but still included in the total count since their nonviability was a pre-existing property of the seedstock. From the germination counts, days to 50% germination (T_50_) was calculated as a measure of germination time, using the “PROBIT” procedure in SAS 9.4 software, which calculates a maximum likelihood estimate of germination timing with a default maximum iteration of 50 (University Edition; SAS Institute, 2017). In the case of germination increasing from 0 to 90-100% between only two time points, model convergence was not attained, and thus the T_50_ value estimated was that of the last maximum likelihood iteration, which was approximately a linear midpoint between the two time points (this occurred for 37 / 225 data points). Negative T_50_ values were excluded (1 / 225). At least two replicate dishes had to have at least five viable, imbibed seeds and at least one germinant in the allotted time in order to be included for germination T_50_ analyses; most accessions exceeded this. Seeds remaining un-imbibed after four weeks were re-scarified precisely using a scalpel and placed with the scar-side down onto the water-saturated sand substrate to determine if their non-germination was due to insufficient scarification; these seeds were then checked weekly for two to three weeks. The final total of germinated seeds was used to calculate germination proportion by accession. Seeds which were damaged from scarification but still germinated were included in germination proportion but not germination T_50_ due to potential changes in germination rate; seeds which were severely damaged from scarification were excluded for both traits. Accession age (years in frozen storage) was the only covariate for germination T_50_ and proportion. Any ungerminated, imbibed seeds by the end of the experiment that were not severely decayed were tested for viability by bisecting the seeds and treating them with tetrazolium chloride (TZ) overnight (Gosling, 2009; Patil and Dadlani, 2009). Dormancy (if present) was broken by physical scarification for most accessions; only two accessions had >5 imbibed, still viable, ungerminated seeds after all germination tests: W6 2427 (*Lathyrus aphaca*: 84% viable ungerminated; 0% germinated) and PI 494749 (*Vicia dumetorum*: 22% viable ungerminated; 25% germinated). This viability was considered evidence that some physiological requirements for germination were not completely met in these accessions; thus, their data for germination T_50_ and proportion was dropped from further analyses.

### Vegetative growth measurements

A subset of seedlings from at least two petri dish replicates of each accession was planted in Ball Professional Growing Mix (no peat; West Chicago, Illinois, USA) within 38-cell trays (cell diameter: 1.95”, cell depth: 4.98”) as soon as possible following germination. For each accession, 4 - 38 seedlings (mean: 15.3 +/- 5.6; after removal of any problematic data) were transplanted and measured and were arranged in replicate groups across trays (1-7 replicates per accession, mean: 3.6); since a subset of perennial seedlings were retained for year two growth, usually more replicates were planted for perennial than annual accessions. From June 19, 2017 - August 25, 2017, plants grew in hoophouse A-6 at the Missouri Botanical Garden (St. Louis, Missouri, USA), which was covered in a light shade cloth; day-night temperatures ranged from 14 - 40°C. Plants were watered typically daily, and 150 ppm 15-5-15 NPK aqueous fertilizer was applied approximately weekly, beginning the earliest week that any plants were measured (for day 21 growth; see below for details); days from most recent fertilization ranged from one to nine days, with the majority measured within five days. Exceptions included some replicates of nine accessions, where their first measurements were taken after less than a full day had elapsed from the most recent fertilization, and one accession that had to be measured after transport to a different greenhouse (see below), which was measured 13 days after the most recent fertilization (at 35 days after planting). We tested our models with these plants dropped respectively to determine if this influenced the results (statistical analysis section). Beginning at the first measurement (∼21 days after planting), climbing plants were trained around thin bamboo poles for taller plants (usually *Phaseolus*) or 18-inch hyacinth sticks for shorter plants. Growth trays were spatially randomized in the hoophouse on August 1, 2017, to avoid potential microclimate heterogeneity. Pesticides were applied on August 15, 2017 (after the majority of plants had been measured) in the hoophouse to control thrips, consisting of a liquid cocktail of Pylon (3mL/gal), Mavrik (3mL/gal), and _Tristar (_ ⅛ teaspoon/quart). A subset of plants had to be transferred on August 25, 2017 to the Saint Louis University (SLU) Biology Department greenhouse (see growth measurements below), where 13 accessions of a mix of annual and perennial *Lathyrus* and *Vicia* had their second measurement taken over the course of two weeks after transfer (up to September 5); here plants were watered every few days, temperatures were similar to the previous hoophouse, but there was somewhat heterogeneous lighting. Due to any impact of transfer or environmental differences, linear models were also tested with these accessions dropped (statistical analysis section).

Vegetative growth was assessed non-destructively for each individual plant over a two-week period. Planting date in the hoophouse was used as a baseline, since seedlings began to show most growth development after this date. All seedlings were first assessed for height, leaf number, vigor, and reproductive status beginning 20-22 days after planting (DAP-21; time point one) in the hoophouse, with the same measurements taken at 34-37 days after planting (DAP-35; time point two) to determine growth rate on an individual plant level, as well as the singular height and leaf measurements at the respective time points (“static” growth measurements). Stem height was measured from ground level to the base of the shoot apex on the tallest stem. Leaf number was counted for each node on the same stem measured for height, up to the most recently developed countable node, at least ∼2 mm from the next node near the shoot apex. If a potential node was leafless, a leaf was only counted if there was a clear remaining structure indicating a leaf was present, such as stipules. For all *Phaseolus* species in this study, two unifoliate eophylls are present at the first true node and thus two leaves were counted for that node for all individuals, even if they had dehisced. Otherwise, there were only single, alternate leaves present along the stem for all genera, with some variation in the number of leaflets per leaf. Lastly, the leaves of *Lathyrus aphaca* are reduced to tendrils, and the stipules are enlarged, foliaceous, and are functionally the main photosynthetic organ of the plant; thus each pair of stipules was counted as one leaf (Sharma and Kumar, 2012). Absolute growth rate (AGR) was calculated on a per-day basis by dividing the difference in height and leaf number by the number of days elapsed between the two time points (as defined in Rees et al., 2010; Pommerening and Muszta, 2016). Relative growth rate (RGR) was calculated for height and leaf number by taking the difference of the natural log of the trait at both dates and dividing by the days elapsed: (ln(trait DAP-35) – ln(trait DAP-21)/(time point 2 – time point 1) (as in Perez-Harguindeguy et al., 2013). Leaf number RGR is also known as relative leaf production rate (RLPR; Garnier, 1992; Grotkopp et al., 2002).

Plants with severe damage were removed from analysis; this included large cuts, bends, or extensive dead tissue in the stem being measured which could have compromised growth, or otherwise irreparable damage to the whole plant. If the damage occurred by DAP-21, all vegetative data was excluded; if the damage occurred by DAP-35, only DAP-35, AGR, and RGR data were removed. Any accessions were dropped for a trait if fewer than three total plants remained following removal of problematic data. A vigor covariate was assessed for each plant qualitatively at both static time points, which considered the impacts of environmental and endogenous factors; this was scored categorically as “low” (high damage and/or was unhealthy, with a severe impact on growth), “medium” (average growth, with moderate to high damage or generally unhealthy, but with a nonlethal impact on growth), or “high” (little to no damage and an overall robust form, with effectively unimpeded growth). The lowest vigor observed between the two dates was assigned as the covariate for AGR and RGR analyses. Due to the possible preferential allocation of energy to reproductive structures, we scored the reproductive status of the plant (“reproductive” or “nonreproductive”) at both dates measured (if either date was “reproductive”, that was assigned to AGR and RGR as well). Lastly, for AGR and RGR, height at DAP-21 was included as a covariate to help control for the effect of different developmental sizes at the start of the growth period. Each of these covariates were tested as random effects in our linear models (statistical analysis section).

### Statistical analyses

Statistical analyses were performed in R v. 3.6.1 (R Core Team, 2019). All analyses were performed on the dataset following removal of any problematic data. Principal component analyses (PCAs) were computed on data scaled to unit variance using the “prcomp” function (base R). PCA was first implemented for accession-level data for all traits in the full dataset (Fig. 1) and then for individual-level seed and vegetative datasets for each genus separately (Figs. S1-S3). In the case of individual-level vegetative data, individuals were dropped if measurement data was missing from an entire date (DAP-21 or DAP-35; and thus growth rate was missing as well). For both accession and individual-level PCAs, any remaining missing data values were imputed using a regularized iterative PCA algorithm (“imputePCA” function in the “missMDA” package; Josse and Husson, 2016), for which the number of components were estimated using generalized cross-validation (“estim_ncpPCA” function). PCs were retained which had an eigenvalue of 1 or greater; in each case these PCs cumulatively explained at least >80% of the total variation in the dataset. Eigenvalues were derived using the “get_eigenvalue” function from the “factoextra” package (Kassambara and Mundt, 2020). PCA figures were created using the package “ggplot2” (Wickham, 2016), and the variable PCA plot was created using the package “ggbiplot” (Vu, 2011). For individual-level genus seed and vegetative PCAs (Fig. S1-S3), PC axes were rotated so they could all be visualized in the same orientation. Accessions and individuals with much reduced data (about half of the traits missing) or which had removed outliers (see below) were excluded from PCAs. Based on these criteria, six accessions were dropped from the accession-level PCA, one each of *Lathyrus annuus*, *L. aphaca*, *L. cicera*, *Phaseolus acutifolius*, *Vicia dumetorum*, and *V. hirsuta*, and imputation was necessary for 7 out of 1,258 data values. For individual-level *Lathyrus* vegetative data, 44 out of 370 individuals were dropped and six out of 2608 data values were imputed. For *Phaseolus*, 50 out of 474 individuals were dropped and no imputation was necessary. For *Vicia*, 36 out of 353 individuals were dropped and one out of 2536 data values was imputed. There were no missing values for individual seed data of any genus.

**Figure 1.**
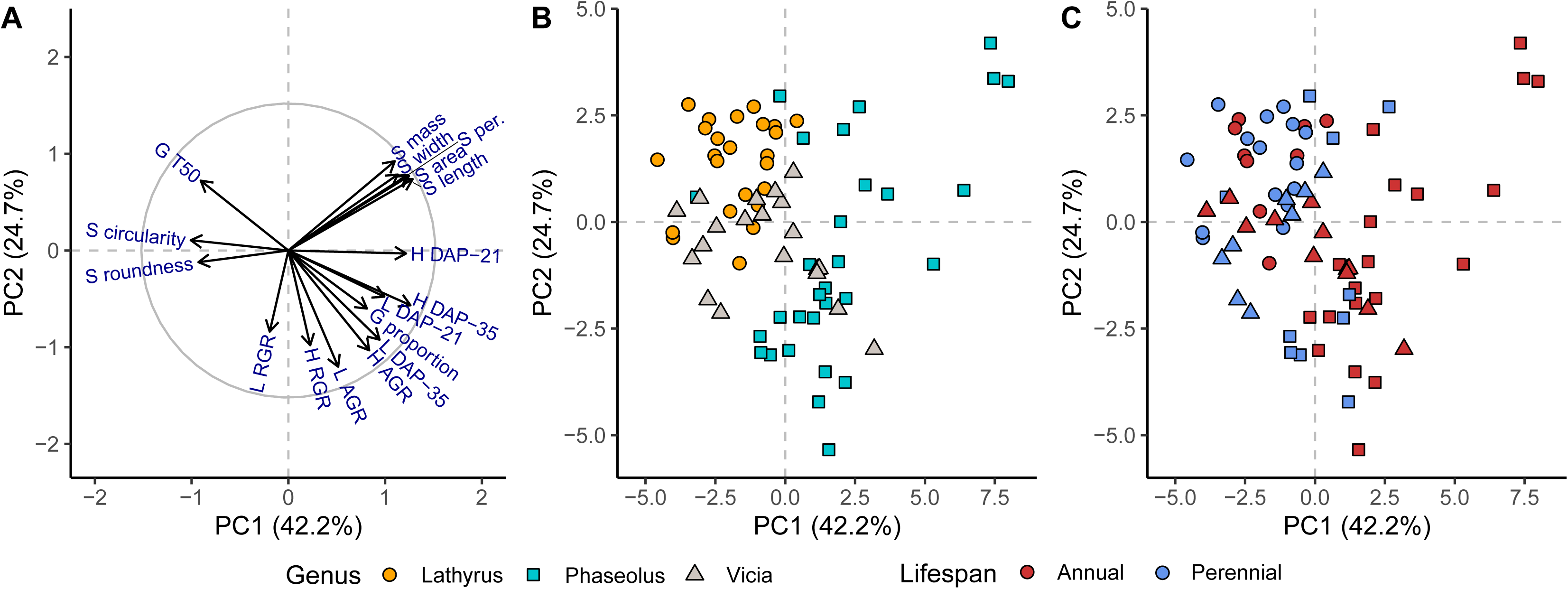
Principal component analysis for the full dataset accession means. (A) shows the relative contribution of each variable on the first two principal components along with the correlation circle; distance of the arrow from the origin indicates increasing representation of that trait in the PCA in a particular region of PC space. Label abbreviations: S signifies seed (perim. is perimeter), G signifies germination, H signifies height, and L signifies leaf. AGR is absolute growth rate, and RGR is relative growth rate. Variable labels were sometimes adjusted slightly from arrow tips to allow complete visualization. (B) shows the individual accession data points in the same PC space colored by genus and (C) the individual accession data points colored by life span.

Linear mixed models were employed on a trait-by-trait basis to assess the contribution of genus, life span, species, and covariates on phenotypic variation using the “lmer” function in the R package “lme4” (Bates et al., 2015). For all ImageJ seed traits (all seed traits excluding seed mass) and all vegetative traits, models were tested on individual seed- and plant-level data respectively. For germination T_50_, the model was tested on individual replicate-level data. The remaining trait models used accession-level data. Each mixed model included genus, life span, genus × life span, and species (nested within genus × life span) as the main fixed effects and accession nested within species as a random effect (except for PC1, PC2, seed mass, and germination proportion, since they measured at the accession level). For growth traits, replicate nested within accession was additionally included in the model as a random effect. Trait models were assessed using a type III ANOVA with Satterthwaite’s method for degrees of freedom, with the exception of PC, seed mass, and germination proportion models, which were assessed using a type I ANOVA (and the base R “lm” function) due to the data consisting of only accession-level means with no significant random effects. Type III ANOVA significance was assessed using the package “afex” (Singmann et al., 2020) along with “lme4.” Using the R package “emmeans,” adjusted means for each genus-life span combination were derived from our models (“emmeans” function) and post hoc custom contrasts were conducted using a Bonferroni correction (“contrast” function; Lenth, 2020). The mentioned covariates under germination and vegetative measurements were added as random effects for the appropriate trait model. The significance of random effects in our models was assessed by taking the likelihood ratio test (LRT) of the model following sequential removal of each random term, using the “rand” function in the package “lmerTest” (Kuznetsova et al., 2017); nonsignificant random effects were dropped from the final model (the “reduced model”). In some cases, certain accessions or species were dropped from models (mentioned above) in order to assess their influence where potential confounding factors were present.

Trait-by-trait correlations were calculated in order to identify phenotypic relationships in the full dataset and within each genus and life span group. Correlation matrices were generated from accession-level data using Pearson correlations (“cor” function in base R) on scaled and centered data, with p values being assessed at a 95% confidence level (“cor.mtest” function in the “corrplot” package; Wei and Simko, 2017). Correlation networks were created from this correlation data using the package “igraph”, with nodes representing traits (organized with “tkplot” function; Csardi and Nepusz, 2006). Nonsignificant correlations (Pearson; *P* < 0.05) were excluded from correlation networks. Full correlation matrix plots for each data group were also generated from Pearson correlation coefficient values using the “corrplot” function (“corrplot” package).

Accessions with trait values which were statistical outliers in their genus, defined as being greater than 1.5 × the interquartile range below or above the 25% and 75% quartiles, respectively, were removed from all analyses of that trait (models, PCA, and correlations) if their sampling replication was also reduced. This was done to reduce the impact of outliers in statistical modelling and estimation of correlations/networks. This resulted in the removal of one accession of *Phaseolus acutifolius* (PI 535200) and *Vicia dumetorum* (PI 494749) for germination T_50_ (also removed due to incomplete germination), and one accession of *Lathyrus annuus* (PI 358829) for height DAP-21 and height and leaf number AGR and RGR (which were dependent on height DAP-21 as a covariate).

## RESULTS

We investigated variation and correlation in seed, germination, and vegetative growth traits of plants derived from germinated seeds in 80 accessions of annual and perennial *Lathyrus*, *Phaseolus*, and *Vicia* species. Principal component analyses highlighted distinct phenotypic space occupied by each genus and life span group, as well as important species-level variation within each genus. Linear models identified genus as the primary predictor of seed size variation and life span as the primary predictor of static vegetative growth variation (the singular height and leaf measurements at DAP-21 and DAP-35). Correlation networks further demonstrate genus- and life span-specific trait relationships, including correlations between and within seed and vegetative traits.

### Genus and life span predict trait variation

Principal component analyses identified key phenotypic differences in the dataset among genera and between life spans, and showed correlated groups of traits (Fig. 1). The first two principal components explained 42.2 and 24.7% of the variance respectively. Seed size traits clustered tightly (Fig. 1A) and loaded positively on both PC1 and PC2. Seed shape (circularity and roundness) loaded negatively onto PC1 (Fig. 1A; Appendix S2), suggesting that seed size and seed shape are negatively correlated. Vegetative traits were less tightly grouped: height at DAP-21 and DAP-35 and leaf number at DAP-21 loaded positively onto PC1, while leaf number at DAP-35 and both growth rate metrics (AGR and RGR) loaded negatively onto PC2 (Fig. 1A; Appendix S2). Germination T_50_ loaded positively onto PC2, opposite of vegetative traits, whereas germination proportion loaded positively onto PC1 in conjunction with vegetative traits (Fig. 1A; Appendix S2). *Phaseolus* occupied the greatest area in the first two PCs, followed by *Vicia* then *Lathyrus* (Fig. 1B). Annuals occupied a greater PC area than perennials, with much of this area occupied by *Phaseolus* annuals (Fig. 1C). Among genera, *Phaseolus* accessions tended to occupy space showing greater seed size and greater static plant height and leaf number, as well as higher growth rate; *Lathyrus* and *Vicia* tended to have smaller, more circular seeds with a delayed T_50_ and smaller vegetative growth. Likewise, annual accessions showed greater seed size and vegetative traits (static and growth rate), while perennials showed the opposite pattern (Fig. 1B,C).

Principal component analyses of seed and vegetative data highlighted similar seed trait correlations in each genus (Appendix S3, S4, S5); however, differences among genera existed in vegetative trait correlations and in how annual and perennial species separated in phenotypic space. In each of the three genera, seed size traits were tightly correlated and loaded positively onto PC1, while seed shape (circularity and roundness) loaded positively onto PC2, approximately orthogonal to seed size (Appendix S3, S4, S5, S6). These data consistently supported a negative relationship between seed size and shape. In general, annual and perennial species did not separate consistently in PC space across genera with respect to seed size; however, *Phaseolus* annuals had more distinctly less circular / round seeds than perennials (Appendix S3, S4, S5). In contrast, vegetative traits showed less consistent and less closely grouped loading patterns in PCAs across genera (Appendix S3, S4, S5, S7). Similar to seed PCAs, *Lathyrus* and *Vicia* had more overlap in annual-perennial vegetative variation than *Phaseolus*. Annual *Phaseolus* species had consistently higher static vegetative growth than perennial species (Appendix S3, S4, S5). Nevertheless, two annual *Vicia* species, *V. villosa* and *V. benghalensis*, showed distinctly greater static vegetative growth than other *Vicia* species (Appendix S5). These data revealed important species-level variation underlying the broad lifespan patterns within each genus.

In our linear mixed models, genus was the largest predictor of seed trait variation (Table 2). Genus was significant for all seed traits except seed mass, whereas life span was not significant for any seed trait (Table 2). Despite life span nonsignificance, post hoc tests revealed that for *Phaseolus* all seed size traits except seed mass were significantly larger in annuals (Appendix S8). For seed circularity and roundness there was a significant genus × life span interaction where both traits were significantly greater in perennials than annuals for *Phaseolus*, but roundness was significantly lower in perennials than annuals for *Lathyrus* (Table 2 and Appendix S8). Linear models revealed a significant genus and life span effect for germination T_50_, while only genus was significant for germination proportion (Table 2). Perennials in all genera had delayed T_50_ compared to annuals, but this was only significant for *Lathyrus* (Appendix S8).

**Table 2.** Analysis of variance (ANOVA) table of the final. reduced linear mixed models for the full dataset. including accession-level principal components (Figure 1) and seed. germination. and vegetative growth traits.

|  | Trait | Genus | Life span | Genus × Life span | Species | Accession |
| --- | --- | --- | --- | --- | --- | --- |
| (a) | PC1 | $F_{2,47} = \mathbf{58.61^{***}}$ | $F_{1,47} = \mathbf{24.89^{***}}$ | $F_{2,47} = \mathbf{5.33^{**}}$ | $F_{21,47} = \mathbf{4.33^{***}}$ | NA<sup>a</sup> |
| | PC2 | $F_{2,47} = \mathbf{24.39^{***}}$ | $F_{1,47} = 0.02$ | $F_{2,47} = 0.36$ | $F_{21,47} = \mathbf{5.98^{***}}$ | NA<sup>a</sup> |
| (b) | Seed mass | $F_{2,49} = 2.04$ | $F_{1,49} = 3.49$ | $F_{2,49} = 0.95$ | $F_{23,49} = \mathbf{5.38^{***}}$ | NA<sup>a</sup> |
| (c) | Seed length | $F_{2,50.8} = \mathbf{21.80^{***}}$ | $F_{1,50.7} = 0.71$ | $F_{2,50.8} = 0.41$ | $F_{23,50.7} = \mathbf{7.84^{***}}$ | LRT = $\mathbf{4708.51^{***}}$ |
| | Seed width | $F_{2,50.8} = \mathbf{15.01^{***}}$ | $F_{1,50.7} = 0.27$ | $F_{2,50.8} = 1.81$ | $F_{23,50.8} = \mathbf{10.24^{***}}$ | LRT = $\mathbf{4029.91^{***}}$ |
| | Seed perimeter | $F_{2,50.8} = \mathbf{21.00^{***}}$ | $F_{1,50.7} = 0.51$ | $F_{2,50.8} = 0.19$ | $F_{23,50.8} = \mathbf{8.92^{***}}$ | LRT = $\mathbf{4737.46^{***}}$ |
| | Seed area | $F_{2,50.8} = \mathbf{19.09^{***}}$ | $F_{1,50.7} = 0.41$ | $F_{2,50.8} = 0.14$ | $F_{23,50.7} = \mathbf{6.93^{***}}$ | LRT = $\mathbf{5113.95^{***}}$ |
| | Seed circularity | $F_{2,49.9} = \mathbf{53.58^{***}}$ | $F_{1,49.2} = 1.07$ | $F_{2,49.9} = \mathbf{6.30^{**}}$ | $F_{23,49.5} = \mathbf{4.97^{***}}$ | LRT = $\mathbf{538.28^{***}}$ |
| | Seed roundness | $F_{2,50.3} = \mathbf{27.43^{***}}$ | $F_{1,49.7} = 0.12$ | $F_{2,50.3} = \mathbf{19.08^{***}}$ | $F_{23,49.9} = \mathbf{4.78^{***}}$ | LRT = $\mathbf{674.03^{***}}$ |
| (d) | Germination T<sub>50</sub> | $F_{2,44.5} = \mathbf{9.07^{***}}$ | $F_{1,44.5} = \mathbf{7.22^*}$ | $F_{2,44.5} = 2.35$ | $F_{20,44.9} = 1.35$ | LRT = $\mathbf{81.64^{***}}$ |
| (e) | Germination proportion | $F_{2,50} = \mathbf{21.62^{***}}$ | $F_{1,50} = 1.44$ | $F_{2,50} = 1.62$ | $F_{22,50} = 0.87$ | NA<sup>a</sup> |
| (f) | Height DAP-21 | $F_{2,47.3} = 1.56$ | $F_{1,47.6} = \mathbf{9.05^{**}}$ | $F_{2,47.3} = 1.63$ | $F_{22,47.1} = 0.85$ | LRT = $\mathbf{222.99^{***}}$ |
| | Leaf number DAP-21 | $F_{2,53.0} = \mathbf{20.83^{***}}$ | $F_{1,60.5} = \mathbf{22.03^{***}}$ | $F_{2,52.9} = 2.70$ | $F_{23,46.9} = \mathbf{6.06^{***}}$ | LRT = $\mathbf{69.38^{***}}$ |
| (g) | Height DAP-35 | $F_{2,49.0} = 1.07$ | $F_{1,49.5} = \mathbf{12.74^{***}}$ | $F_{2,48.9} = 1.73$ | $F_{23,48.1} = \mathbf{1.96^*}$ | LRT = $\mathbf{158.00^{***}}$ |
| | Leaf number DAP-35 | $F_{2,52.0} = \mathbf{17.16^{***}}$ | $F_{1,54.1} = \mathbf{13.19^{***}}$ | $F_{2,51.6} = 3.00$ | $F_{23,48.5} = \mathbf{9.37^{***}}$ | LRT = $\mathbf{41.05^{***}}$ |
| (h) | Height AGR | $F_{2,48.1} = 0.26$ | $F_{1,48.8} = 1.53$ | $F_{2,47.9} = 0.58$ | $F_{22,46.8} = \mathbf{2.51^{**}}$ | LRT = **103.65\*\*\*** |
| | Height RGR | $F_{2,45.5} = 0.33$ | $F_{1,46.9} = 0.17$ | $F_{2,45.4} = 1.10$ | $F_{22,44.0} = \mathbf{2.70^{**}}$ | LRT = **54.46\*\*\*** |
| | Leaf number AGR | $F_{2,49.5} = 1.04$ | $F_{1,51.1} = 0.06$ | $F_{2,49.1} = 0.23$ | $F_{22,46.5} = \mathbf{3.00^{***}}$ | LRT = **43.24\*\*\*** |
| | Leaf number RGR | $F_{2,49.3} = 1.31$ | $F_{1,52.4} = 3.05$ | $F_{2,49.3} = 1.21$ | $F_{22,47.1} = 1.64$ | LRT = **30.27\*\*\*** |

\* $P < 0.05$ ; \*\* $P < 0.01$ ; \*\*\* $P < 0.001$ .
**Notes:** Letters denote separate models with different random effects, while the main effects are the same for all traits. The accession effect and other significant random effects from the reduced model were included; the accession effect is represented by the likelihood ratio test statistic (LRT). ANOVAs are all type III with the exception of PC1, PC2, seed mass, and germination proportion (type I), due to the data consisting of only accession-level means with no significant random effects. Additional random effect significance is listed in Appendix S9. Significant values are bolded (at least $P < 0.05$ ). When non-whole numbers, denominator degrees of freedom were rounded to the first decimal.
<sup>a</sup> Accession could not be used as a random effect in these trait models due to the dataset consisting of accession-level means.

Life span was the most consistent predictor of vegetative trait variation (Table 2). Life span was a significant predictor for all static height and leaf number measurements at DAP-21 and DAP-35, with more variation attributable to life span than genus (nonsignificant) for static height traits (Table 2). For static leaf traits, life span explained a similar amount of variation to genus, and both effects were significant (Table 2). For all genera, perennials had lower mean static vegetative height and leaf number than congeneric annuals; however, the difference was only significant for *Phaseolus* (all static vegetative traits) and for *Vicia* (static leaf number only; Appendix S8). Vegetative growth rate patterns (AGR and RGR) were more variable, and neither genus nor life span was significant for these traits in the linear models (Table 2). However, *Lathyrus* showed a significant difference in leaf number RGR (perennials greater than annuals) and *Phaseolus* in height AGR and RGR (annuals greater than perennials; Appendix S8). Although nonsignificant, in *Lathyrus* both height and leaf AGR and height RGR were also higher in perennials; in *Vicia* height and leaf number AGR were higher in annuals than perennials, but height and leaf number RGR were higher in perennials (nonsignificant; Appendix S8). In addition, species was a significant predictor for most traits, with the exception of germination T_50_, germination proportion, height at DAP-21, and leaf number RGR (Table 2). Several random effects controlled for in the models were significant (Appendix S9), while dropping data with potentially confounding factors from the models resulted in minimal change in significance (Appendix S10).

### Genus and life span patterns in trait correlations

Correlation networks revealed dynamic seed and vegetative trait correlations which reflected unique genus and life span-specific patterns, and which were generally consistent with PCA variable loadings (Fig. 2). Considering the full dataset network and commonalities among each subgroup, there were always strong, significant positive correlations among all seed size traits, which were always significantly negatively correlated with seed circularity and/or roundness (Fig. 2; Appendix S11). Germination T_50_ was usually negatively correlated with at least some vegetative traits for all subgroups, while germination proportion was generally positively correlated with vegetative traits; both germination traits generally positively correlated with seed traits when significant (Fig. 2). All significant correlations among vegetative growth traits were positive, with the exception of height and leaf number RGR, but significance and magnitude of correlation among vegetative traits varied for each subgroup (Fig. 2). Seed size traits were more commonly significantly correlated to static vegetative growth traits (positive) than AGR or RGR (Fig. 2). Where significant correlations did occur, height and leaf number RGR usually negatively correlated with seed size (Fig. 2). Overall, static height measurements had a consistently high degree (number of significant connections to other traits) in each subgroup (Fig. 2). Despite commonalities, considerable variation in magnitude, direction, and significance in trait integration also existed among the subgroups.

**Figure 2.**
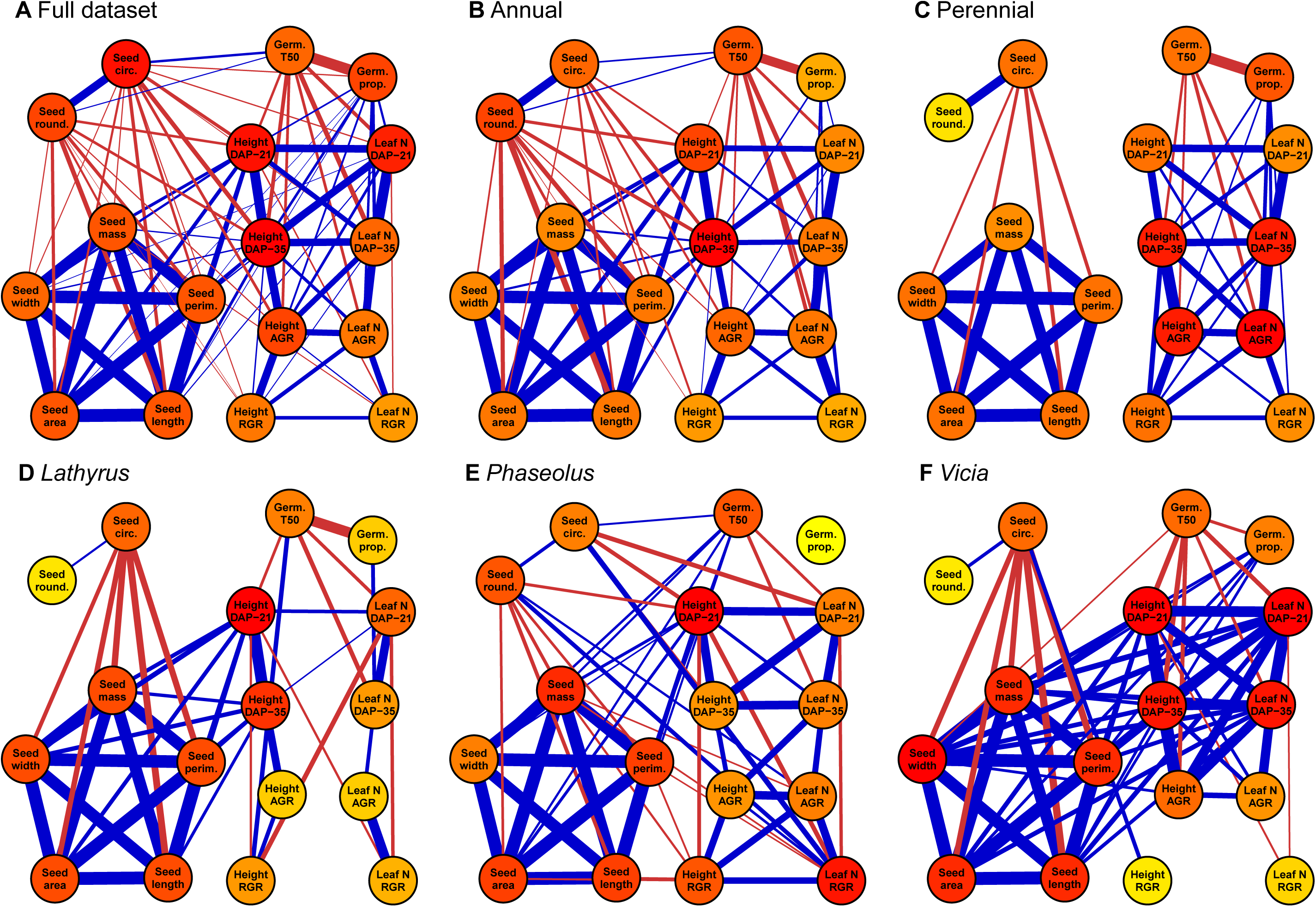
Correlation networks for (A) the full dataset, and the dataset subgroups: (B) annuals, (C) perennials, (D) *Lathyrus*, (E) *Phaseolus*, and (F) *Vicia*. Presence of lines (edges) between trait nodes indicates a significant correlation between those traits (Pearson; *P* < 0.05). Blue signifies a positive correlation and red a negative correlation; line thickness corresponds to the strength of the correlation. Node color signifies degree (the number of significant trait connections to that node), which ranges from yellow (low) to red (high); note that color is relative to the maximum number of connections for that subgroup and so is not directly comparable across subgroups.

The three genera differed in the extent of integration between seed and vegetative traits, as well as the strength of correlations among vegetative traits (Fig. 2D-E). *Lathyrus* showed the least amount of significant seed to vegetative trait correlation, with only static height positively correlated to seed size (Fig. 2D; Appendix S12). *Lathyrus* also showed the least connectivity among vegetative traits, with height and leaf trait groups having few significant correlations (Fig. 2D, Appendix S12). *Phaseolus* showed significant positive correlations between seed size and height at DAP-21 and significant negative correlations between seed size and height/leaf RGR, with the opposite pattern for seed shape traits with vegetative traits (Fig. 2E, Appendix S13). *Phaseolus* also showed predominantly positive correlations between vegetative traits, and it had more significant seed and vegetative trait connections to RGR than the other genera (Fig. 2E, Appendix S13). *Vicia* showed the greatest amount of connectivity between seed and vegetative traits, with all static DAP-21 and DAP-35 height and leaf measurements significantly positively correlated with all seed size traits, although there was minimal correlation between AGR/RGR and seed traits (Fig. 2F, Appendix S14). *Vicia* also showed the most robust cluster of significant positively correlated vegetative traits, with the exception of RGR (Fig. 2F, Appendix S14). Lastly, seed size was significantly negatively correlated to seed circularity for *Lathyrus* and *Vicia*, while for *Phaseolus* seed size traits were significantly negatively correlated with seed roundness but not circularity (Fig. 2D-F, Appendix S12, S13, S14). For all three genera, height at DAP-21 had a consistently high degree (Fig. 2D-F).

Annual and perennial species differed in their seed trait to vegetative growth correlations and their overall connectivity among traits (Fig. 2B-C). While annuals showed significant positive relationships between all seed size and static height measurements, as well as significant negative correlations between seed shape and all height traits, perennials lacked any significant correlation between seed and vegetative traits (Fig. 2B-C; Appendix S15, S16). Annuals also had more significant correlations between seed size and seed shape traits (negative) than perennials, and annuals showed significant positive correlations between seed shape and germination T_50_, while seed and germination traits were not connected for perennials (Fig. 2B-C; Appendix S15, S16). However, both annual and perennial subgroups showed significant positive correlations among most vegetative traits, with the only gaps in correlation occurring for height and leaf AGR/RGR (Fig. 2B-C; Appendix S15, S16). The degree of seed and static vegetative traits was low in perennials compared to annuals, but perennials generally had greater positive correlations between height and leaf traits, as well as between static height and height AGR/RGR (Fig. 2B-C; Appendix S15, S16). Similar to the genera subgroups, static vegetative traits showed some of the highest degrees within both the annual and perennial networks (Fig. 2B-C).

## DISCUSSION

Life history strategy and the associated life span classification (annual, perennial) is associated with various aspects of species reproductive and vegetative biology, but the role of life span in predicting seed to vegetative trait correlation is not well characterized. Here we examined trait variation and correlation of seed size and shape, and the germination and vegetative growth characters of seedlings derived from those seeds, in annual and perennial congeners from three herbaceous legume genera. We found that while genus was a stronger predictor of seed size and shape, life span most consistently predicted static vegetative growth. Patterns of trait correlation between and among seed size and vegetative growth traits differed by life span and genus; specifically, annual species had more significant seed to vegetative trait correlations than perennial species.

### Evolutionary basis of genus and life span differences

In this study genus was consistently the greatest predictor of seed size and shape variation, which illustrates the importance of phylogenetic context in understanding some aspects of life history variation (Silvertown and Dodd, 1996). Annual-perennial overlap in species-level variation presented here in PCAs further demonstrates that seed size and shape variation does not consistently separate by life span group (Appendix S3, S4, S5). Mazer (1989) similarly found a greater amount of variation in seed mass explained by phylogenetic family (30%) than life history type (22%) in a large study of ten families of Indiana Dune angiosperms. However, they still found life history to be significant, which was attributed to the distinctly large seeds of tree species (Mazer, 1989). Our study builds on this previous work in that it focuses on herbaceous species of three legume genera and quantifies multiple dimensions of seed size and shape on an individual seed level (instead of seed mass class), without specificity to a particular habitat (Mazer, 1989). Importantly, Mazer (1989) also noted that life history was usually not a significant predictor of seed mass looking at the within-family level; also, there was not a significant difference in seed mass among annuals and herbaceous perennials in their study. Our lack of clear life span signal in seed size (but see post hoc tests for *Phaseolus*) is also consistent with a meta-analysis of c. 3000 congeneric comparisons of annual and perennial species (Vico et al., 2016).

While genus predicted seed traits, our data demonstrated that life span consistently predicted static vegetative traits. In this study, annual species displayed greater mean height and leaf number measured at 21 and 35 days from planting than congeneric perennial species in the first year of growth; this was significant for both traits in *Phaseolus* and for leaf number in *Vicia*. Higher growth in annuals is consistent with the predicted pattern for annuals and perennials according to the acquisitive-conservative resource economics spectrum (e.g., Roumet et al., 2006, González-Paleo and Ravetta, 2015). Vegetative growth is commonly measured in terms of relative growth rate (RGR), for which life span is usually a significant predictor (e.g., Garnier, 1992; Atkinson et al., 2016); however, in this study neither life span nor genus was a significant predictor of either absolute growth rate (AGR) or RGR in terms of plant height and leaf number. This could reflect the timing of growth measurements (made at the seedling stage), since species have different trajectories of RGR over their life span, with RGR typically decreasing with time as the plant becomes larger (Turnbull et al., 2008). Also, while RGR is typically measured using successive destructive biomass harvests on different plants (Perez-Harguindeguy et al., 2013), we nondestructively measured height and leaf number growth on the same individual plants, which may result in different patterns. Ideally, vegetative measurements will span a much larger developmental window, with multiple time points allowing for more precise modeling of growth through the life cycle of each plant.

Annual species exhibited significant correlations between seed and vegetative traits (plant height and leaf number), but the same correlations were not significant in perennial species, suggesting that some phenotypic relationships are at least partially dependent on life span (Fig. 2). Other studies of herbaceous perennial species have focused on relationships between vegetative traits and traits of seeds produced by that individual or from individuals in separate studies, a slightly different approach than our study, where we examined correlations between seed traits and the vegetative traits expressed by the individuals germinated from those seeds. Kleyer et al. (2019), studying predominantly herbaceous perennials in a NW European flora, also found no significant correlation between plant height and mass of seeds produced by that plant, although seed mass was somewhat correlated to plant biomass. Similarly, a study of 526 species in Sweden found no significant relationship between plant height and seed mass in herbaceous perennials, while that relationship did exist for herbaceous annuals, woody perennials, and the full dataset (Bolmgren and Cowen, 2008). In this case seed and plant height traits were combined from separate studies in a meta-analysis (Bolmgren and Cowen, 2008). Our study extends these results and demonstrates similar correlation patterns between source seeds and the plants derived from them. Significant seed to vegetative trait correlations in annual species could be related to annuals’ complete reliance on seeds to perpetuate their population; thus many aspects of annual plant form should be optimized for producing a greater reproductive output. Plant size and seed size are often significantly positively correlated in broad interspecific meta-analyses (Leishman et al., 1995; Moles et al., 2004; Díaz et al., 2016); greater vegetative size may allow greater total reproductive output through increased seed size and/or number.

Unique properties of this dataset must also be taken into consideration for their influence on correlation trends. For example, life span seed vs. vegetative trends were heavily influenced by *Phaseolus* accessions, which tended to have some of the highest trait values but were nonetheless supported by multiple replicated accessions. Further research with greater replication will be necessary to determine if this intergeneric life span trend is consistent within each genus. Phenotypic integration is also sensitive to growth environment and developmental stage (Murren, 2002); thus it remains to be seen if these phenotypic patterns are robust in each species’ native environment and across a longer period of growth, particularly in the case of perennial species.

This study examined a broad diversity of annual and perennial interspecific variation, but variation at the intraspecific level was not as well represented. The fact that species was highly significant for almost all traits reflects the plethora of phenotypic diversity that exists within and among annual and perennial species. For simplicity, we adopted the broad titles of annual and perennial in this study, but life span is not a simple binary or categorical trait, and life span alone cannot be expected to encompass the biological complexity of all underlying traits. There is substantial intra- and interspecific variation in traits associated with life span: reproductive patterns (e.g., semelparous and iteroparous), growth determinacy vs. indeterminacy, clonality, length of the juvenile phase, and mating system (autogamous, allogamous, and mixed), among many others (Friedman, 2020). Life span is also inherently context dependent and potentially variable at the intraspecific level: both annual and perennial populations can occur within a single species across its range, often due to variation in environmental disturbance, e.g., *Mimulus guttatus* (Friedman et al., 2015), *Oryza perennis* (Morishima et al., 1984), and *Zostera marina* (Reynolds et al., 2017). We are also limited in our ability to know and control for the maternal growth environment and past environmental drivers for these accessions, which may include either direct or indirect artificial selection for those collected in feral populations and as agricultural weeds. In order to gain a more precise understanding of life span variation in these species, a thorough assessment of variation within and across many diverse populations is needed.

### Relevance to perennial breeding goals

With increasing interest in *de novo* domestication of wild species and breeding of hybrid herbaceous perennial crops, broadening our understanding of trait networks in herbaceous perennials is essential. Much of our understanding of phenotypic change and correlation in domestication targeting reproductive structures comes from annual and woody perennial cultivars (Miller and Gross, 2011; Meyer et al., 2012), and a gap exists in the case of herbaceous perennials, which have not been broadly domesticated for human grain consumption (Van Tassel et al., 2010). There is evidence that annual crops exhibit a decrease in phenotypic integration during crop improvement, potentially releasing them from ecological trait trade-offs present in more resource-limited wild conditions (Milla et al., 2014). Further characterization of functional traits such as those in this study can advance understanding of evolution under domestication and can also help in identifying promising wild candidates for domestication. Evidence to date highlights potential seed yield - vegetative trade-offs in emergent perennial crops (e.g., González-Paleo et al., 2016; Pastor-Pastor et al., 2018); however, the challenge of co-selecting for negatively correlated traits has been accomplished in modern plant breeding, and breeders have successfully selected for both high seed yield and sustained perenniality in rice (DeHaan et al., 2005; Huang et al., 2018).

Seed size is a key trait in perennial grain breeding programs (Kantar et al., 2016), and the effect of seed size selection on other plant phenotypes is relevant to understanding constraints to evolution under artificial selection. Our results suggest that different phenotypic ties exist between seed and early vegetative growth traits in annuals and perennials measured under controlled conditions, with seed traits ostensibly being less integrated with growth traits in perennials under these circumstances. In these genera, this could mean that selection on seed size may not greatly impact early vegetative growth in perennials. This could be particularly important for grain crops which are dual-purposed as forage, where vigorous vegetative growth is also favorable, e.g., *Thinopyrum intermedium* (Pugliese, 2017) and *Silphium integrifolium* (Vilela et al., 2020). It remains to be seen if the seed-vegetative disconnect extends to later lifetime vegetative characteristics such as biomass allocation and plant size at reproductive maturity, and if these relationships hold when mature plants are measured under field conditions.

The next steps in investigating phenotypic integration in herbaceous perennials should extend to traits important to perennial agriculture and total perennial lifetime fitness, measured in the field over multiple years. Specifically, total seed yield and root allocation (among other belowground and perennating structures) are dual targets of current perennial grain breeding, and understanding potential trade-offs between these traits is fundamental to the advancement of perennial breeding (Van Tassel et al., 2017). Our results highlight phenotypic patterns only in the first few weeks of perennial growth, whereas multiyear patterns in yield and vegetative allocation in perennial species will more comprehensively reflect their total life history strategy. While logistically difficult, understanding yearly fluctuations in reproductive output, above-ground biomass, and root growth over the total life span of perennials is paramount, as will be connecting this variation to environmental variables. By gathering this lifetime phenotypic information, early seed and growth traits may be used to predict hard-to-measure later life span aspects of perennial crops and thus expedite the breeding process.

## CONCLUSIONS

Despite the prevalence of annual-perennial transitions in angiosperms, we know relatively little about how life history classification is associated with phenotypic correlations across ecologically and agriculturally important traits. Here we show that in three genera of legumes seed variation was primarily explained by genus and static vegetative variation by life span. Further, annual species showed stronger seed to vegetative trait correlation in early growth than perennial congeners. These findings call for further investigation into how these trait correlations differ throughout the life span of perennial plants, as well as into correlations among other important life history traits, particularly allocation to reproductive and perennating organs. This study highlights that both life span and phylogenetic context are important in predicting phenotypic variation, and there remains numerous underexplored systems in which to expand life history studies.

## ACKNOWLEDGEMENTS

This research was funded by the Perennial Agriculture Project (Malone Family Land Preservation Foundation and The Land Institute). S.A.H. was supported by a graduate assistantship from Saint Louis University. M.J.R. is supported by the Donald Danforth Plant Science Center and the Perennial Agriculture Project. The National Science Foundation Research Experiences for Undergraduates (REU) program at the Missouri Botanical Garden supported student involvement in this research, including M.C.S. A special thanks also goes to REU student Summer Sherrod for her assistance in this work and enthusiastic engagement in this project as a part of her internship. Seeds were provided by the United States Department of Agriculture Western Regional PI Station (Pullman, WA); from the USDA facility in Pullman, we specifically thank curators Clarice Coyne and Barbara Hellier for their assistance in clarifying improvement status and other provenance details for each accession. We are grateful to Daniel Debouck (former curator of *Phaseolus* at the International Center for Tropical Agriculture) for invaluable discussions regarding appropriate species life span assignment. We are grateful to the greenhouse and horticulture staff at the Missouri Botanical Garden for providing space and assistance in plant care, especially Joshua Higgins, Justin Lee, and Derek Lyle. We would also like to thank the Saint Louis University Department of Biology and Donald Danforth Plant Science Center, and specifically Kasey Fowler-Finn, Kristina Haines, and Kevin Reilly, for assistance with plant growth facilities. We acknowledge the following researchers who helped provide important information regarding accession origin: Alexandr Afonin, Clarice Coyne, Ken Friesen, Stephanie Greene, Richard Hannan, Barbara Hellier, Douglas Johnson, Francis Kilkenny, Nigel Maxted, Luis Guillermo Santos Meléndez, Sergey Shuvalov, and Filip Vandelook. We are thankful to the Miller Lab and Elizabeth Kellogg for helpful comments on previous versions of this manuscript, and specifically for Zachary Harris and Julia Pratt for assistance in figure generation. Lastly, we are especially grateful to all who assisted in plant measurement and caretaking: Niyati Bhakta, Emily Boeckenstedt, Leah Brand, Claudia Ciotir, Emma Frawley, Jordan Hathaway, Danielle Hopkins, Tanvi Kadiyala, Aidan Leckie-Harre, Alex Linan, Kazi Maharun Nessa, Brittany Pace, Joshua Reinl, Heather Schier, and William Shoenberger.

## AUTHOR CONTRIBUTIONS

S.A.H. and A.J.M. designed the study, and S.A.H. implemented the research and wrote the manuscript. M.A.A., Q.G.L., A.J.M., M.J.R., and M.C.S. assisted in critical review and writing of the manuscript. M.J.R. assisted in germination calculations and statistical methods and interpretation. M.A.A. and Q.G.L. assisted in crafting germination experiments and provided the necessary resources. M.C.S. assisted in data acquisition.

## DATA AVAILABILITY

All phenotypic data (individual seed, germination, vegetative growth, and accession-level data) are available in the Figshare database. Data can be accessed at: doi: [data is currently in preparation for submission to this database].

## SUPPORTING INFORMATION

Additional supporting information may be found online in the Supporting Information section at the end of the article.

## APPENDIX S1.

Accession descriptive metadata and sample size for each trait.

## APPENDIX S2.

Variable loadings for principal components 1-5 of the full accession-level dataset principal component analysis (Fig. 1).

## APPENDIX S3.

Principal component analyses of the full seed and vegetative trait datasets for *Lathyrus*.

## APPENDIX S4.

Principal component analyses of the full seed and vegetative trait datasets for *Phaseolus*.

## APPENDIX S5.

Principal component analyses of the full seed and vegetative trait datasets for *Vicia*.

## APPENDIX S6.

Variable loadings for principal components 1-2 of the full seed dataset principal component analysis, parsed by genus.

## APPENDIX S7.

Variable loadings for the full vegetative dataset principal component analysis, parsed by genus (principal components 1-3 for *Lathyrus*, principal components 1-2 for *Phaseolus* and *Vicia*).

## APPENDIX S8.

Table of adjusted means of each trait for annuals and perennials of each genus and post hoc significance tests.

## APPENDIX S9.

Table of all significant random effects in addition to accession for each trait model.

## APPENDIX S10.

Analysis of variance table of each trait model with subsets of the data removed to test for influence on significance.

## APPENDIX S11.

Correlation matrix for the full dataset, showing Pearson correlation coefficients between every combination of traits, using accession-level data.

## APPENDIX S12.

Correlation matrix for the *Lathyrus* data subgroup, showing Pearson correlation coefficients between every combination of traits, using accession-level data.

## APPENDIX S13.

Correlation matrix for the *Phaseolus* data subgroup, showing Pearson correlation coefficients between every combination of traits, using accession-level data.

## APPENDIX S14.

Correlation matrix for the *Vicia* data subgroup, showing Pearson correlation coefficients between every combination of traits, using accession-level data.

## APPENDIX S15.

Correlation matrix for the annual data subgroup, showing Pearson correlation coefficients between every combination of traits, using accession-level data.

## APPENDIX S16.

Correlation matrix for the perennial data subgroup, showing Pearson correlation coefficients between every combination of traits, using accession-level data.

