## Appendix S2 for "The role of genus and life span in predicting seed and vegetative trait variation and correlation in *Lathyrus*, *Phaseolus*, and *Vicia* (Fabaceae)"

**Appendix S2.** Variable loadings for each principal component used for the full dataset accession-level PCA. The first five PCs are reported since their eigenvalue was greater than 1; cumulatively they explain > 90% of the variation in the dataset. Important loadings are bolded, defined as having a loading value that is greater than what it would be if all variables equally contributed to the PC’s variation (square root of one divided by the number of variables (17); that is, greater than 0.24253563). Variables are organized according to the standard order of traits in this study (e.g., Appendix S8) and the variable importance, starting with PC1.

| Variable | PC1 loading | PC2 loading | PC3 loading | PC4 loading | PC5 loading |
| --- | --- | --- | --- | --- | --- |
| Seed mass | **0.27014990** | **0.29655555** | -0.08035681 | 0.19310018 | 0.01526868 |
| Seed length | **0.31566175** | 0.23698417 | -0.14260123 | 0.06397929 | 0.01474618 |
| Seed width | **0.27779612** | **0.25220573** | -0.11430082 | **0.28515259** | 0.04846663 |
| Seed perimeter | **0.30616859** | **0.24731207** | -0.14091150 | 0.13620108 | 0.02260461 |
| Seed area | **0.30296046** | **0.25036488** | -0.13973899 | 0.14074329 | 0.01867071 |
| Seed circularity | **-0.24783049** | 0.03414804 | 0.16395916 | **0.52143508** | 0.06768769 |
| Height DAP-21 | **0.29881535** | -0.00996964 | **0.32063217** | -0.11123683 | 0.12832846 |
| Height DAP-35 | **0.30998107** | -0.18187903 | 0.13160487 | -0.12036061 | 0.23149426 |
| Leaf number DAP-21 | **0.24360593** | -0.15329629 | **0.45323965** | 0.03803732 | 0.14105835 |
| Height AGR | 0.20527423 | **-0.33145057** | -0.14707176 | -0.09824012 | **0.27496535** |
| Height RGR | 0.05547198 | **-0.31462917** | **-0.43512117** | -0.11782441 | 0.22037716 |
| Leaf number DAP-35 | 0.23158743 | **-0.29626930** | **0.24838650** | 0.20736394 | 0.13231836 |
| Leaf AGR | 0.12645058 | **-0.38585560** | -0.12509867 | **0.34535400** | 0.09054718 |
| Leaf RGR | -0.04735759 | **-0.26945141** | **-0.45649411** | **0.33181492** | -0.05812795 |
| Seed roundness | -0.22825753 | -0.03876145 | **0.25008288** | **0.48383995** | 0.09573648 |
| Germination T_50_ | -0.22260650 | 0.23197326 | -0.09163646 | 0.01228113 | **0.54775598** |
| Germination proportion | 0.19950953 | -0.19278325 | 0.02635743 | 0.09268175 | **-0.66283033** |
