## Appendix S5 for "The role of genus and life span in predicting seed and vegetative trait variation and correlation in *Lathyrus*, *Phaseolus*, and *Vicia* (Fabaceae)"

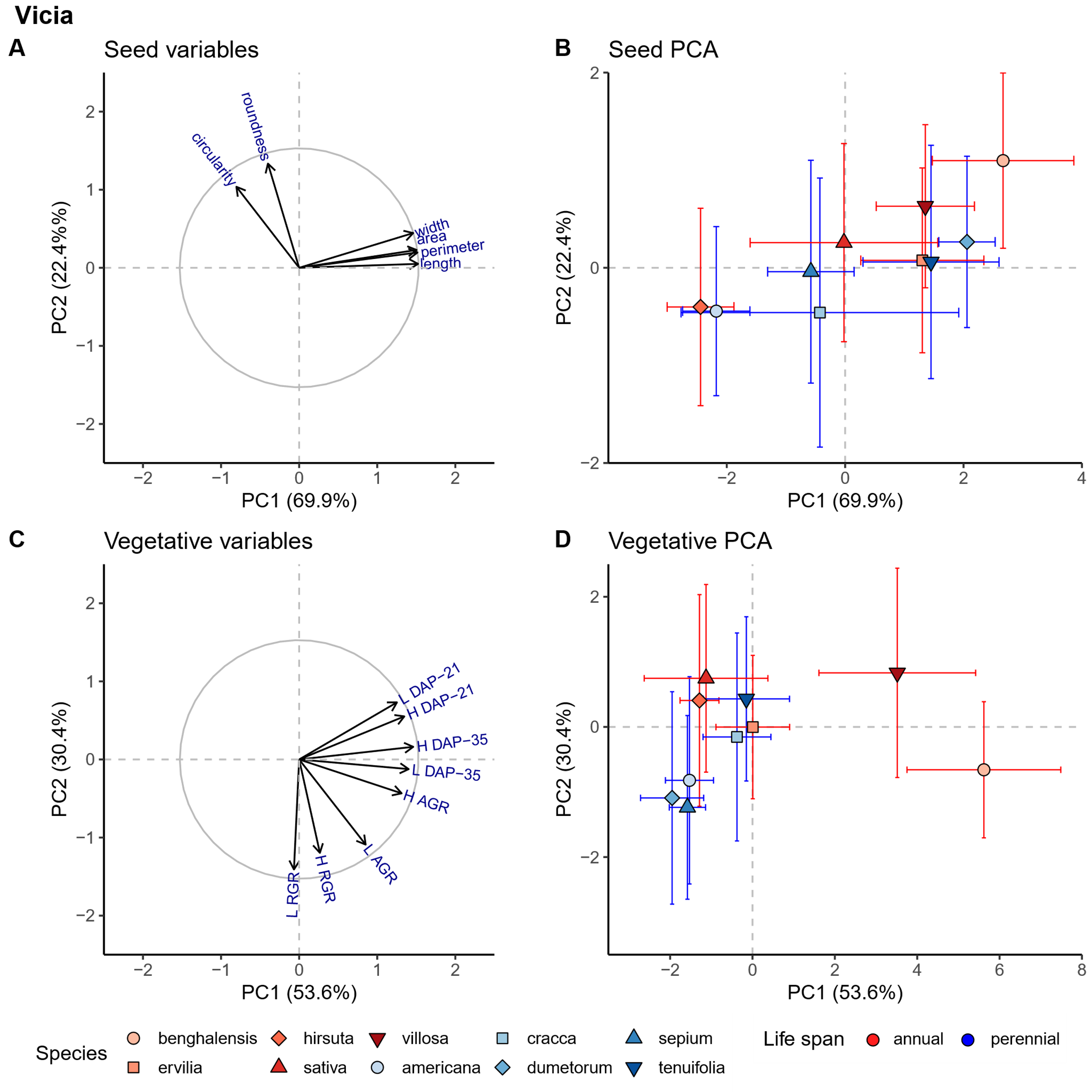

**Appendix S5.** Principal Component Analyses of the full individual-level seed and vegetative trait dataset for *Vicia*. Variable plots (A,C) show the variable loadings onto the seed and vegetative trait PCAs respectively; distance of the arrow from the origin indicates increasing representation of that trait in the PCA in a particular region of PC space. Each species is represented in PC space (B,D), where central points represent the species mean and error bars represent one standard deviation. Different species have different shapes and color shades, with red signifying annual species and blue signifying perennial species.
