## Appendix S6 for "The role of genus and life span in predicting seed and vegetative trait variation and correlation in *Lathyrus*, *Phaseolus*, and *Vicia* (Fabaceae)"

**Appendix S6.** Variable loadings for each principal component used for the full seed dataset PCA, parsed by genus. The first two PCs for each genus are reported on the grounds that their eigenvalue is greater than 1; cumulatively they explain > 90% of the variation in each genus’s dataset. Important loadings are bolded, defined as having a loading value that is greater than what it would be if all variables equally contributed to the PC’s variation (square root of one divided by the number of variables (6); that is, greater than 0.40824829). Variables are organized according to the standard order of traits in this study (e.g., Appendix S8) and the variable importance, starting with PC1.

| Genus | Variable | PC1 loading | PC2 loading |
| --- | --- | --- | --- |
| *Lathyrus* | Seed length | **0.48775246** | -0.03741388 |
|  | Seed width | **0.46211994** | 0.27968674 |
|  | Seed perimeter | **0.48901022** | 0.06699677 |
|  | Seed area | **0.48390306** | 0.10020232 |
|  | Seed circularity | -0.26852835 | **0.55446050** |
|  | Seed roundness | -0.05605404 | **0.77357612** |
| Genus | Variable | PC1 loading | PC2 loading |
| *Phaseolus* | Seed length | **0.49688105** | -0.02601732 |
|  | Seed width | **0.47610305** | 0.21785655 |
|  | Seed perimeter | **0.49677460** | 0.05416632 |
|  | Seed area | **0.49269097** | 0.06903796 |
|  | Seed circularity | -0.04238172 | **0.72637200** |
|  | Seed roundness | -0.18737529 | **0.64540306** |
| Genus | Variable | PC1 loading | PC2 loading |
| *Vicia* | Seed length | **0.4859405** | 0.02834453 |
|  | Seed width | **0.4661613** | 0.25081730 |
|  | Seed perimeter | **0.4837607** | 0.11058372 |
|  | Seed area | **0.4800846** | 0.12866882 |
|  | Seed circularity | -0.2564292 | **0.58487483** |
|  | Seed roundness | -0.1276474 | **0.75194699** |
