## Appendix S7 for "The role of genus and life span in predicting seed and vegetative trait variation and correlation in *Lathyrus*, *Phaseolus*, and *Vicia* (Fabaceae)"

**Appendix S7.** Variable loadings for each principal component used for the full vegetative dataset PCA, parsed by genus. The number PCs reported is based on the grounds that their eigenvalue is greater than 1; cumulatively they explain > 80% of the variation in each genus’s dataset. Important loadings are bolded, defined as having a loading value that is greater than what it would be if all variables equally contributed to the PC’s variation (square root of one divided by the number of variables (8); that is, greater than 0.35355339). Variables are organized according to the standard order of traits in this study (e.g., Appendix S8) and the variable importance, starting with PC1.

| Genus | Variable | PC1 loading | PC2 loading | PC 3 loading |
| --- | --- | --- | --- | --- |
| *Lathyrus* | Height DAP-21 | **0.42833300** | 0.31943403 | -0.01374738 |
|  | Height DAP-35 | **0.52766926** | 0.07631637 | -0.29737122 |
|  | Height AGR | **0.38820486** | -0.24931140 | **-0.49517959** |
|  | Leaf number DAP-21 | **0.35857215** | 0.33409279 | 0.22958404 |
|  | Leaf number DAP-35 | **0.44842691** | -0.11994184 | **0.46390774** |
|  | Height RGR | 0.05957556 | **-0.43174440** | **-0.44586007** |
|  | Leaf AGR | 0.23190797 | **-0.47319674** | **0.37210328** |
|  | Leaf RGR | 0.02004371 | **-0.54190324** | 0.24699648 |
| Genus | Variable | PC1 loading | PC2 loading |  |
| *Phaseolus* | Height DAP-35 | **0.4308963** | 0.25165338 |  |
|  | Height AGR | **0.4431014** | -0.15550982 |  |
|  | Leaf number DAP-35 | **0.4757147** | 0.09915621 |  |
|  | Leaf AGR | **0.4119739** | -0.27858333 |  |
|  | Height DAP-21 | 0.1902060 | **0.49402476** |  |
|  | Height RGR | 0.2370515 | **-0.42484189** |  |
|  | Leaf number DAP-21 | 0.3024181 | **0.42050288** |  |
|  | Leaf RGR | 0.1952795 | **-0.47294067** |  |
| Genus | Variable | PC1 loading | PC2 loading |  |
| *Vicia* | Height DAP-21 | **0.42577111** | 0.23164002 |  |
|  | Height DAP-35 | **0.46183392** | 0.06783053 |  |
|  | Height AGR | **0.41491327** | -0.18218681 |  |
|  | Leaf number DAP-21 | **0.39591628** | 0.30705589 |  |
|  | Leaf number DAP-35 | **0.44339350** | -0.05189188 |  |
|  | Height RGR | 0.08396809 | **-0.50214484** |  |
|  | Leaf AGR | 0.26913690 | **-0.45775903** |  |
|  | Leaf RGR | -0.02103734 | **-0.59150742** |  |
