## Appendix S8 for "The role of genus and life span in predicting seed and vegetative trait variation and correlation in *Lathyrus*, *Phaseolus*, and *Vicia* (Fabaceae)"

**Appendix S8.** Adjusted trait means with one standard error, derived from reduced linear models, grouped by genus and life span. Letters show the results of post hoc custom contrasts for each model, with covariates included: different letters denote a significant difference (at least *P* < 0.05) for a trait between life span groups within a genus (bolded), with a Bonferroni correction applied to the p-value. AGR signifies absolute growth rate and RGR relative growth rate. PC1 and PC2 are the first two principal components of the full dataset accession-level PCA. Values less than one are rounded to two significant figures (with the exception of principal components).

| Genus |  | Life span | |  |
| --- | --- | --- | --- | --- |
| *Lathyrus* | Trait | Annual | Perennial | Significance |
|  | PC1 | -1.23 ± 0.50 ^A^ | -1.95 ± 0.40 ^A^ | NS |
|  | PC2 | 1.43 ± 0.44 ^A^ | 1.05 ± 0.35 ^A^ | NS |
|  | Seed mass (mg) | 41.28 ± 5.91 ^A^ | 33.28 ± 5.09 ^A^ | NS |
|  | Seed length (mm) | 4.46 ± 0.25 ^A^ | 4.22 ± 0.22 ^A^ | NS |
|  | Seed width (mm) | 3.82 ± 0.16 ^A^ | 3.54 ± 0.14 ^A^ | NS |
|  | Seed perimeter (mm) | 13.63 ± 0.70 ^A^ | 12.75 ± 0.60 ^A^ | NS |
|  | Seed area (mm^2^) | 13.36 ± 1.68 ^A^ | 11.94 ± 1.45 ^A^ | NS |
|  | Seed circularity | 0.88 ± 0.0029 ^A^ | 0.89 ± 0.0025 ^A^ | NS |
|  | Seed roundness | **0.89 ± 0.011 ^A^** | **0.86 ± 0.0093 ^B^** | * |
|  | Germination T_50_ (d) | **9.54 ± 1.63 ^A^** | **17.73 ± 1.15 ^B^** | *** |
|  | Germination proportion | **0.86 ± 0.049 ^A^** | **0.67 ± 0.041 ^B^** | * |
|  | Height DAP-21 (mm) | 88.98 ± 41.65 ^A^ | 64.64 ± 34.85 ^A^ | NS |
|  | Height DAP-35 (mm) | 176.41 ± 60.85 ^A^ | 124.42 ± 57.36 ^A^ | NS |
|  | Height AGR (mm/d) | 4.03 ± 3.01 ^A^ | 4.77 ± 2.80 ^A^ | NS |
|  | Height RGR (d **^-1^**) ^a^ | 0.019 ± 0.0062 ^A^ | 0.029 ± 0.0048 ^A^ | NS |
|  | Leaf number DAP-21 | 5.81 ± 0.79 ^A^ | 4.79 ± 0.75 ^A^ | NS |
|  | Leaf number DAP-35 | 8.90 ± 1.18 ^A^ | 8.36 ± 1.15 ^A^ | NS |
|  | Leaf number AGR (leaves/d) | 0.17 ± 0.053 ^A^ | 0.25 ± 0.049 ^A^ | NS |
|  | Leaf number RGR (d **^-1^**) ^a^ | **0.018 ± 0.0033 ^A^** | **0.033 ± 0.0025 ^B^** | ** |
| *Phaseolus* | Trait | Annual | Perennial |  |
|  | PC1 | **3.31 ± 0.34 ^A^** | **-0.09 ± 0.48 ^B^** | *** |
|  | PC2 | -0.71 ± 0.29 ^A^ | 0.02 ± 0.42 ^A^ | NS |
|  | Seed mass (mg) | 50.37 ± 4.27 ^A^ | 34.21 ± 6.24 ^A^ | NS |
|  | Seed length (mm) | **5.87 ± 0.18 ^A^** | **4.77 ± 0.26 ^B^** | ** |
|  | Seed width (mm) | **4.36 ± 0.12 ^A^** | **3.86 ± 0.17 ^B^** | * |
|  | Seed perimeter (mm) | **17.06 ± 0.51 ^A^** | **14.31 ± 0.74 ^B^** | ** |
|  | Seed area (mm^2^) | **21.78 ± 1.22 ^A^** | **15.65 ± 1.78 ^B^** | * |
|  | Seed circularity | **0.84 ± 0.0022 ^A^** | **0.86 ± 0.0032 ^B^** | *** |
|  | Seed roundness | **0.77 ± 0.0080 ^A^** | **0.81 ± 0.012 ^B^** | * |
|  | Germination T_50_ (d) | 1.73 ± 1.07 ^A^ | 2.35 ± 1.55 ^A^ | NS |
|  | Germination proportion | 0.96 ± 0.035 ^A^ | 0.96 ± 0.051 ^A^ | NS |
|  | Height DAP-21 (mm) | **243.31 ± 31.39 ^A^** | **68.62 ± 39.86 ^B^** | *** |
|  | Height DAP-35 (mm) | **493.85 ± 54.46 ^A^** | **204.34 ± 60.98 ^B^** | *** |
|  | Height AGR (mm/d) | **16.65 ± 2.69 ^A^** | **8.83 ± 2.93 ^B^** | *** |
|  | Height RGR (d **^-1^**) ^a^ | **0.056 ± 0.0039 ^A^** | **0.039 ± 0.0056 ^B^** | * |
|  | Leaf number DAP-21 | **7.45 ± 0.74 ^A^** | **4.24 ± 0.77 ^B^** | *** |
|  | Leaf number DAP-35 | **12.57 ± 1.13 ^A^** | **9.03 ± 1.17 ^B^** | *** |
|  | Leaf number AGR (leaves/d) | 0.33 ± 0.047 ^A^ | 0.29 ± 0.051 ^A^ | NS |
|  | Leaf number RGR (d **^-1^**) ^a^ | 0.030 ± 0.0021 ^A^ | 0.037 ± 0.0029 ^A^ | NS |
| *Vicia* | Trait | Annual | Perennial |  |
|  | PC1 | -0.33 ± 0.46 ^A^ | -2.08 ± 0.56 ^A^ | NS |
|  | PC2 | -0.83 ± 0.40 ^A^ | -0.39 ± 0.49 ^A^ | NS |
|  | Seed mass (mg) | 32.99 ± 5.99 ^A^ | 26.75 ± 6.70 ^A^ | NS |
|  | Seed length (mm) | 3.92 ± 0.23 ^A^ | 3.75 ± 0.28 ^A^ | NS |
|  | Seed width (mm) | 3.54 ± 0.15 ^A^ | 3.27 ± 0.19 ^A^ | NS |
|  | Seed perimeter (mm) | 12.12 ± 0.64 ^A^ | 11.45 ± 0.79 ^A^ | NS |
|  | Seed area (mm^2^) | 11.15 ± 1.53 ^A^ | 9.92 ± 1.91 ^A^ | NS |
|  | Seed circularity | 0.90 ± 0.0027 ^A^ | 0.90 ± 0.0033 ^A^ | NS |
|  | Seed roundness | 0.93 ± 0.0099 ^A^ | 0.89 ± 0.012 ^A^ | NS |
|  | Germination T_50_ (d) | 3.81± 1.38 ^A^ | 7.88 ± 1.76 ^A^ | NS |
|  | Germination proportion | 0.94 ± 0.044 ^A^ | 0.93 ± 0.058 ^A^ | NS |
|  | Height DAP-21 (mm) | 167.27 ± 38.65 ^A^ | 74.88 ± 42.08 ^A^ | NS |
|  | Height DAP-35 (mm) | 270.10 ± 59.59 ^A^ | 141.49 ± 62.83 ^A^ | NS |
|  | Height AGR (mm/d) | 6.77 ± 2.87 ^A^ | 4.92 ± 2.99 ^A^ | NS |
|  | Height RGR (d **^-1^**) ^a^ | 0.025 ± 0.0055 ^A^ | 0.028 ± 0.0059 ^A^ | NS |
|  | Leaf number DAP-21 | **7.86 ± 0.77 ^A^** | **4.78 ± 0.79 ^B^** | *** |
|  | Leaf number DAP-35 | **12.79 ± 1.16 ^A^** | **8.94 ± 1.18 ^B^** | *** |
|  | Leaf number AGR (leaves/d) | 0.33 ± 0.050 ^A^ | 0.29 ± 0.052 ^A^ | NS |
|  | Leaf number RGR (d **^-1^**) ^a^ | 0.029 ± 0.0030 ^A^ | 0.039 ± 0.0031 ^A^ | NS |

* *P* < 0.05; ***P* < 0.01; ****P* < 0.001; NS = not significant (*P* > 0.05).

^a^ Relative growth rate is calculated as [ln(trait2) – ln(trait1) / time2 – time1], which results in a unitless numerator.
