## Appendix S9 for "The role of genus and life span in predicting seed and vegetative trait variation and correlation in *Lathyrus*, *Phaseolus*, and *Vicia* (Fabaceae)"

**Appendix S9.** Table of all significant random effects in addition to accession for each trait model (accession significance is in Table 2). Significance was evaluated by removing terms and comparison of models. Letters denote models with different random effects. Note: seed size and shape traits are not included here since they had no random effects other than accession. For germination proportion and T_50_, the age covariate was not significant. PC1, PC2, and seed mass were taken at the accession level and also have no random effects. REML is restricted maximum likelihood, used for model fitting. AIC is the Akaike information criterion. LRT is the likelihood ratio test statistic.

|  | **Trait** | **Random effect** | **Parameters** | **REML log-likelihood** | **AIC** | **LRT** |
| --- | --- | --- | --- | --- | --- | --- |
| (a) | Height DAP-21 | Replicate | 31 | -6384.42 | 12830.85 | 14.85*** |
|  |  | Vigor | 31 | -6406.82 | 12875.63 | 59.64*** |
|  | Leaf number DAP-21 | Replicate | 32 | -1961.26 | 3986.52 | 23.13*** |
|  |  | Vigor | 32 | -2029.52 | 4123.05 | 159.66*** |
| (b) | Height DAP-35 | Replicate | 33 | -6404.47 | 12874.93 | 44.26*** |
|  |  | Vigor | 33 | -6425.05 | 12916.11 | 85.43*** |
|  |  | Reproductive status | 33 | -6389.24 | 12844.49 | 13.81*** |
|  | Leaf number DAP-35 | Replicate | 33 | -2333.96 | 4733.91 | 20.99*** |
|  |  | Vigor | 33 | -2359.91 | 4785.81 | 72.90*** |
|  |  | Reproductive status | 33 | -2330.26 | 4726.51 | 13.60*** |
| (c) | Height AGR | Replicate | 32 | -3194.40 | 6452.79 | 21.72*** |
|  |  | Vigor | 32 | -3197.97 | 6459.94 | 28.87*** |
|  |  | Reproductive status | 32 | -3191.56 | 6447.11 | 16.04*** |
|  | Height RGR | Replicate | 31 | 2417.60 | -4773.20 | 23.83*** |
|  |  | Height DAP-21 | 31 | 2422.50 | -4782.99 | 14.04*** |
|  | Leaf AGR | Replicate | 32 | 625.37 | -1186.74 | 5.53* |
|  |  | Vigor | 32 | 623.19 | -1182.38 | 9.89** |
|  |  | Reproductive status | 32 | 624.14 | -1184.27 | 8.00** |
|  | Leaf RGR | Replicate | 30 | 2770.29 | -5480.58 | 5.82* |

* *P* < 0.05; ***P* < 0.01; ****P* < 0.001.
