## Appendix S10 for "The role of genus and life span in predicting seed and vegetative trait variation and correlation in *Lathyrus*, *Phaseolus*, and *Vicia* (Fabaceae)"

**Appendix S10.** Analysis of variance (ANOVA) table of linear mixed models for seed, germination, and vegetative growth traits where data was removed that may influence results in unknown ways. All fixed and significant random effects from the original models are still included, following the same model reduction steps (Table 2). The accession effect is represented by the likelihood ratio test statistic (LRT). The data being removed for each model are: (a) plants measured less than one full day after the first fertilization treatment (only occurred for DAP-21); (b) plants measured 13 days after the most recent fertilization (only for DAP-35); (c) plants that had to be measured at DAP-35 after transport to a separate Saint Louis University greenhouse; (d) with *Phaseolus lunatus* removed; and (e) with the accession removed which had half of its seeds damaged and excluded from germination (*Vicia hirsuta*; PI 219631). ANOVAs are all type III with the exception of seed mass and germination proportion (type I), due to the data consisting of only accession-level means with no significant random effects. Significant values are bolded. When non-whole numbers denominator degrees of freedom were rounded to the first decimal. **See the summary below the table for any changes to model significance compared to the original model (Table 2).**

|  | **Trait** | **Genus** | **Life span** | **Genus × Life span** | **Species** | **Accession** |
| --- | --- | --- | --- | --- | --- | --- |
| (a) | Height DAP-21 | *F*_2, 44.3_ = 1.46 | *F*_1, 44.5_ = **8.45**** | *F*_2, 44.3_ = 1.55 | *F*_22, 44.3_ = 0.75 | LRT = **240.00***** |
|  | Leaf number DAP-21 | *F*_2, 48.8_ = **20.16***** | *F*_1, 55.6_ = **21.62***** | *F*_2, 48.8_ = 2.72 | *F*_23, 43.3_ = **6.00***** | LRT = **58.61***** |
|  | Height AGR | *F*_2, 45.2_ = 0.29 | *F*_1, 45.8_ = 1.84 | *F*_2, 45.0_ = 0.97 | *F*_22, 43.8_ = **3.82***** | LRT = **76.80***** |
|  | Height RGR | *F*_2, 42.1_ = 0.63 | *F*_1, 43.4_ = 0.56 | *F*_2, 42.1_ = 0.92 | *F*_22, 40.9_ = **2.58**** | LRT = **48.09***** |
|  | Leaf number AGR | *F*_2, 48.3_ = 1.06 | *F*_1, 50.0_ = 0.01 | *F*_2, 47.7_ = 0.16 | *F*_22, 43.6_ = **4.13***** | LRT = **24.65***** |
|  | Leaf number RGR | *F*_2, 46.9_ = 1.72 | *F*_1, 49.9_ = 3.76 | *F*_2, 46.9_ = 1.10 | *F*_22, 43.7_ = **1.90*** | LRT = **20.68***** |
| (b) | Height DAP-35 | *F*_2, 48.0_ = 1.04 | *F*_1, 48.5_ = **12.48***** | *F*_2, 47.9_ = 1.69 | *F*_23, 47.1_ = **1.92*** | LRT = **157.34***** |
|  | Leaf number DAP-35 | *F*_2, 51.0_ = **16.99***** | *F*_1, 53.0_ = **13.10***** | *F*_2, 50.6_ = 2.96 | *F*_23, 47.5_ = **9.32***** | LRT = **40.75***** |
|  | Height AGR | *F*_2, 47.0_ = 0.25 | *F*_1, 47.7_ = 1.51 | *F*_2, 46.9_ = 0.57 | *F*_22, 46.2_ = **2.47**** | LRT = **102.77***** |
|  | Height RGR | *F*_2, 44.8_ = 0.34 | *F*_1, 46.1_ = 0.19 | *F*_2, 44.7_ = 1.07 | *F*_22, 43.9_ = **2.68**** | LRT = **54.16***** |
|  | Leaf number AGR | *F*_2, 48.6_ = 1.03 | *F*_1, 50.1_ = 0.06 | *F*_2, 48.2_ = 0.23 | *F*_22, 46.3_ = **2.95***** | LRT = **43.01***** |
|  | Leaf number RGR | *F*_2, 48.4_ = 1.30 | *F*_1, 51.4_ = 3.00 | *F*_2, 48.4_ = 1.20 | *F*_22, 47.0_ = 1.62 | LRT = **30.10***** |
| (c) | Height DAP-35 | *F*_2, 39.0_ = 0.63 | *F*_1, 39.0_ = **10.51**** | *F*_2, 38.9_ = 2.16 | *F*_20, 39.4_ =1.82 | LRT = **136.60***** |
|  | Leaf number DAP-35 | *F*_2, 39.7_ = **17.27***** | *F*_1, 39.7_ = **26.64***** | *F*_2, 39.4_ = 0.86 | *F*_20, 40.1_ = **8.34***** | LRT = **32.84***** |
|  | Height AGR | *F*_2, 37.2_ = 0.13 | *F*_1, 37.0_ = 1.16 | *F*_2, 37.0_ = 0.92 | *F*_19, 38.6_ = **2.45**** | LRT = **90.45***** |
|  | Height RGR | *F*_2, 35.1_ = 0.68 | *F*_1, 34.8_ = 0.15 | *F*_2, 34.8_ = 0.62 | *F*_19, 37.3_ = **2.55**** | LRT = **53.47***** |
|  | Leaf number AGR | *F*_2, 36.4_ = 0.75 | *F*_1, 36.0_ = 0.66 | *F*_2, 35.9_ = 0.06 | *F*_19, 38.3_ = **2.89**** | LRT = **35.99***** |
|  | Leaf number RGR | *F*_2, 33.7_ = 1.39 | *F*_1, 33.8_ = 3.62 | *F*_2, 33.7_ = 1.36 | *F*_19, 37.4_ = **1.97*** | LRT = **19.51***** |
| (d) | Seed mass | *F*_2, 48_ = 1.00 | *F*_1, 48_ = 2.21 | *F*_2, 48_ = 0.47 | *F*_22, 48_ = **5.24***** | NA ^a^ |
|  | Seed length | *F*_2, 49.8_ = **21.48***** | *F*_1, 49.7_ = 0.70 | *F*_2, 49.8_ = 0.40 | *F*_22, 49.7_ = **7.26***** | LRT = **4705.44***** |
|  | Seed width | *F*_2, 49.8_ = **15.03***** | *F*_1, 49.7_ = 0.27 | *F*_2, 49.8_ = 1.81 | *F*_22, 49.8_ = **8.76***** | LRT = **4013.84***** |
|  | Seed perimeter | *F*_2, 49.8_ = **20.73***** | *F*_1, 49.7_ = 0.50 | *F*_2, 49.8_ = 0.19 | *F*_22, 49.7_ = **8.00***** | LRT = **4726.58***** |
|  | Seed area | *F*_2, 49.8_ = **18.97***** | *F*_1, 49.7_ = 0.41 | *F*_2, 49.8_ = 0.14 | *F*_22, 49.7_ = **5.93***** | LRT = **5160.18***** |
|  | Seed circularity | *F*_2, 49.0_ = **52.76***** | *F*_1, 48.3_ = 1.06 | *F*_2, 49.0_ = **6.20**** | *F*_22, 48.3_ = **4.68***** | LRT = **537.94***** |
|  | Seed roundness | *F*_2, 49.4_ = **27.45***** | *F*_1, 48.8_ = 0.12 | *F*_2, 49.4_ = **19.10***** | *F*_22, 48.8_ = **4.81***** | LRT = **665.90***** |
|  | Germination T_50_ | *F*_2, 43.5_ = **8.87***** | *F*_1, 43.5_ = **7.06*** | *F*_2, 43.5_ = 2.30 | *F*_19, 43.9_ = 1.39 | LRT = **78.99***** |
|  | Germination proportion | *F*_2, 49_ = **20.50***** | *F*_1, 49_ = 1.32 | *F*_2, 49_ = 1.66 | *F*_21, 49_ = 0.88 | NA ^a^ |
|  | Height DAP-21 | *F*_2, 46.4_ = 1.55 | *F*_1, 46.7_ = **8.97**** | *F*_2, 46.4_ = 1.63 | *F*_21, 46.2_ = 0.86 | LRT = **224.70***** |
|  | Leaf number DAP-21 | *F*_2, 52.0_ = **20.43***** | *F*_1, 59.2_ = **21.60***** | *F*_2, 51.9_ = 2.65 | *F*_22, 46.0_ = **6.20***** | LRT = **68.40***** |
|  | Height DAP-35 | *F*_2, 48.0_ = 1.05 | *F*_1, 48.5_ = **12.68***** | *F*_2, 47.9_ = 1.74 | *F*_22, 47.2_ = **1.89*** | LRT = **156.62***** |
|  | Leaf number DAP-35 | *F*_2, 50.9_ = **16.98***** | *F*_1, 52.9_ = **13.06***** | *F*_2, 50.5_ = 2.97 | *F*_22, 47.5_ = **9.33***** | LRT = **40.75***** |
|  | Height AGR | *F*_2, 47.0_ = 0.25 | *F*_1, 47.5_ = 1.47 | *F*_2, 46.8_ = 0.57 | *F*_21, 45.9_ = **2.00*** | LRT = **116.56***** |
|  | Height RGR | *F*_2, 44.9_ = 0.38 | *F*_1, 46.2_ = 0.29 | *F*_2, 44.8_ = 1.08 | *F*_21, 43.5_ = **2.58**** | LRT = **58.44***** |
|  | Leaf number AGR | *F*_2, 48.4_ = 1.02 | *F*_1, 49.9_ = 0.06 | *F*_2, 48.0_ = 0.24 | *F*_21, 45.4_ = **2.80**** | LRT = **42.03***** |
|  | Leaf number RGR | *F*_2, 48.3_ = 1.30 | *F*_1, 51.4_ = 3.02 | *F*_2, 48.3_ = 1.20 | *F*_21, 46.0_ = 1.62 | LRT = **29.25***** |
| (e) | Germination T_50_ | *F*_2, 47.7_ = **7.87**** | *F*_1, 39.6_ = **7.35**** | *F*_2, 42.0_ = 2.03 | *F*_20, 30.2_ = 1.56 | LRT = **9.05**** |
|  | Germination proportion | *F*_2, 49_ = **22.96***** | *F*_1, 49_ = 2.02 | *F*_2, 49_ = 1.51 | *F*_22, 49_ = 0.73 | NA ^a^ |

* *P* < 0.05; ***P* < 0.01; ****P* < 0.001.

^a^ Accession could not be used as a random effect in these trait models due to the dataset consisting of accession-level means.

**Summary:** The following are any changes to the significance (i.e., switch from nonsignificant to significant and vice versa) of the main effects in the above new linear models as compared to the original model with the full dataset (Table 2). For model (a), the species effect for leaf number RGR went from nonsignificant in the original model to significant in the new model (*P* < 0.05). For model (c), the species effect for height DAP-35 went from significant in the original model (*P* < 0.05) to nonsignificant in the new model, and the species effect for leaf number RGR went from nonsignificant in the original model to significant in the new model (*P* < 0.05). No other changes in significance occurred.
