## Appendix S14 for "The role of genus and life span in predicting seed and vegetative trait variation and correlation in *Lathyrus*, *Phaseolus*, and *Vicia* (Fabaceae)"

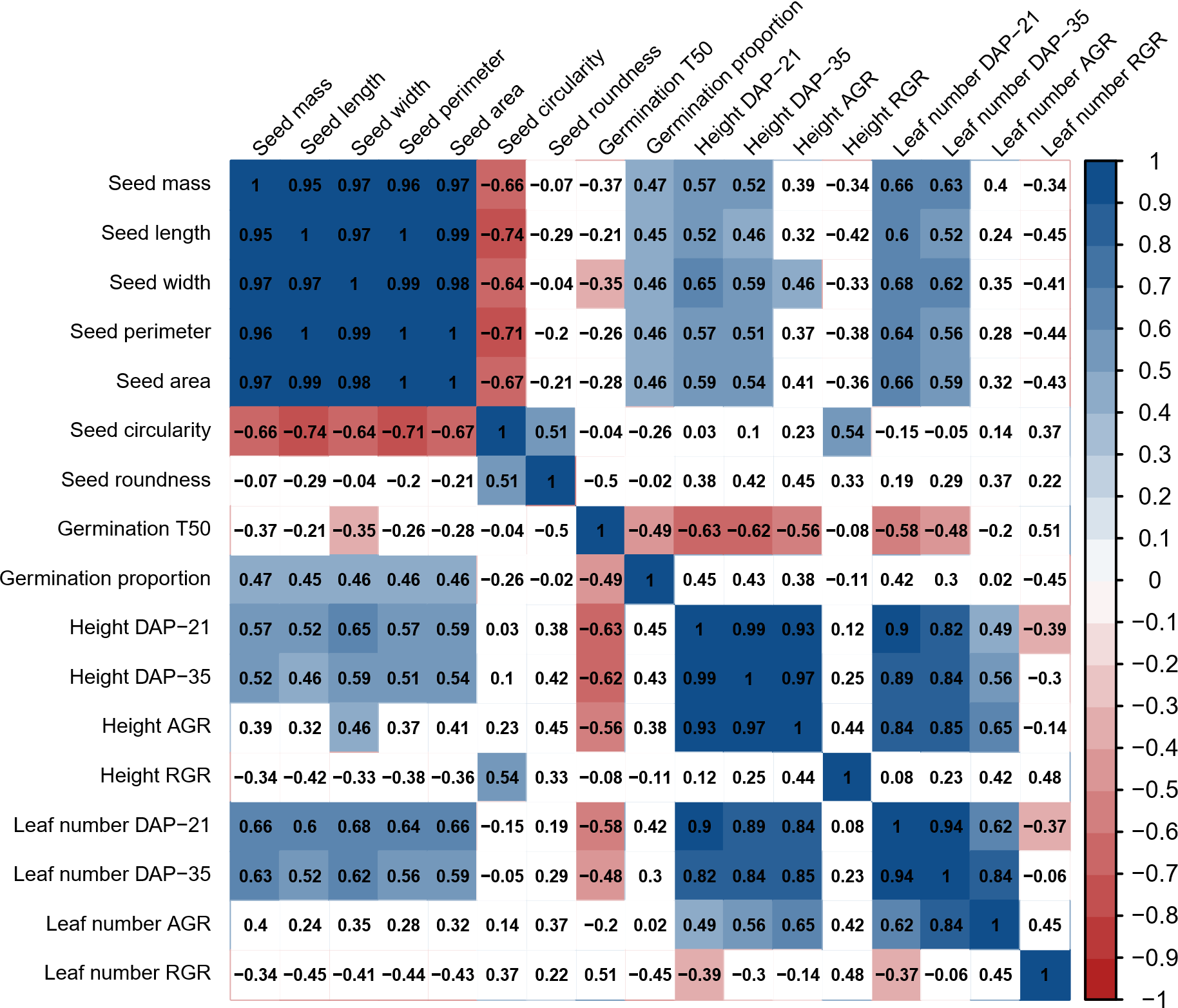

**Appendix S14.** Correlation matrix for the *Vicia* data subgroup, showing Pearson correlation coefficients between every combination of traits at the accession level. Blue and red signify a significant positive and negative correlation, respectively (*P* < 0.05). Squares are white if the correlation is not significant.
