## Appendix S16 for "The role of genus and life span in predicting seed and vegetative trait variation and correlation in *Lathyrus*, *Phaseolus*, and *Vicia* (Fabaceae)"

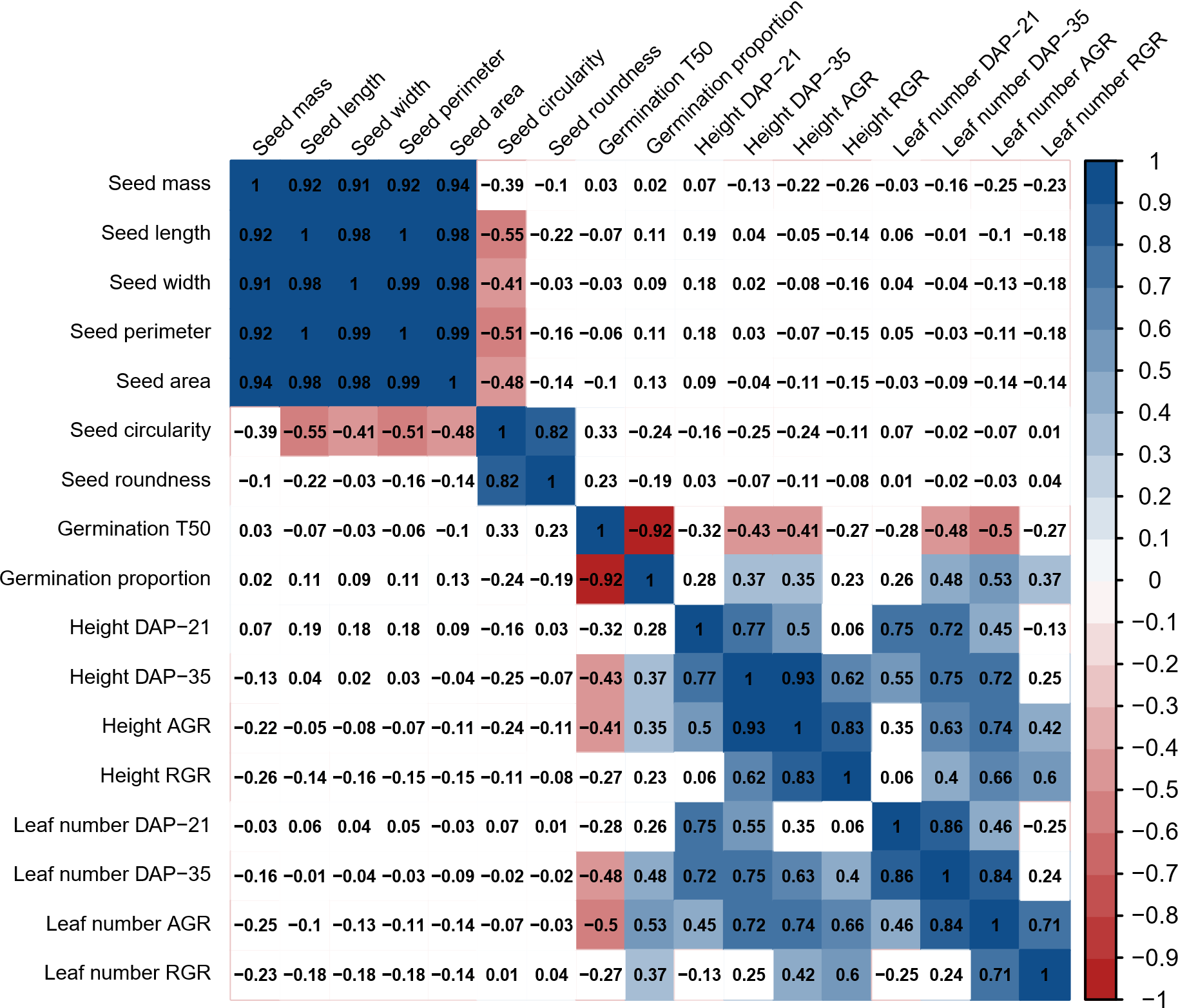

**Appendix S16.** Correlation matrix for the perennial data subgroup, showing Pearson correlation coefficients between every combination of traits, using accession-level data. Blue and red signify a significant positive and negative correlation, respectively (*P* < 0.05). Squares are white if the correlation is not significant.
